# Leprosy in wild chimpanzees

**DOI:** 10.1101/2020.11.10.374371

**Authors:** Kimberley J. Hockings, Benjamin Mubemba, Charlotte Avanzi, Kamilla Pleh, Ariane Düx, Elena Bersacola, Joana Bessa, Marina Ramon, Sonja Metzger, Livia V. Patrono, Jenny E. Jaffe, Andrej Benjak, Camille Bonneaud, Philippe Busso, Emmanuel Couacy-Hymann, Moussa Gado, Sebastien Gagneux, Roch C. Johnson, Mamoudou Kodio, Joshua Lynton-Jenkins, Irina Morozova, Kerstin Mätz-Rensing, Aissa Regalla, Abílio R. Said, Verena J. Schuenemann, Samba O. Sow, John S. Spencer, Markus Ulrich, Hyacinthe Zoubi, Stewart T. Cole, Roman M. Wittig, Sebastien Calvignac-Spencer, Fabian H. Leendertz

**Affiliations:** Centre for Ecology and Conservation, University of Exeter, Penryn TR10 9FE, UK; Centre for Research in Anthropology (CRIA – NOVA FCSH), Lisbon, Portugal; Project Group Epidemiology of Highly Pathogenic Microorganisms, Robert Koch Institute, Berlin, Germany; Department of Wildlife Sciences, School of Natural Resources, Copperbelt University, Kitwe, Zambia; Global Health Institute, Ecole Polytechnique Fédérale de Lausanne, Lausanne, Switzerland; Department of Microbiology, Immunology and Pathology, Colorado State University, Fort Collins, CO, USA; Swiss Tropical and Public Health Institute, Basel, Switzerland; University of Basel, Basel, Switzerland; Taï Chimpanzee Project, Centre Suisse de Recherches Scientifiques, Abidjan, Ivory Coast; Department of Zoology, University of Oxford, Oxford, UK; Department for BioMedical Research, University of Bern, Bern, Switzerland; Laboratoire National d’Appui au Développement Agricole/Laboratoire central de Pathologie Animale, Bingerville, Côte d’Ivoire; Programme National de Lutte contre la Lèpre, Ministry of Public Health, Niamey, Niger; Centre Interfacultaire de Formation et de Recherche en Environnement pour le Développement Durable, University of Abomey-Calavi, 03 BP 1463, Jericho, Cotonou, Benin; Fondation Raoul Follereau, Paris, France; Centre National d’Appui à la Lutte Contre la Maladie, Bamako, Mali; Institute of Evolutionary Medicine, University of Zurich, Winterthurerstrasse 190, 8057, Zurich, Switzerland; Pathology Unit, German Primate Center, Leibniz-Institute for Primate Research, Göttingen; Instituto da Biodiversidade e das Áreas Protegidas, Dr. Alfredo Simão da Silva (IBAP), Av. Dom Settimio Arturro Ferrazzetta, Bissau, Guiné-Bissau; Programme National d’Elimination de la Lèpre, Sénégal; Institut Pasteur, Paris, France; Max Planck Institute for Evolutionary Anthropology, Leipzig, Germany

**Keywords:** *Mycobacterium leprae*, great ape, environmental source, reservoir, whole-genome

## Abstract

Humans are considered the main host for *Mycobacterium leprae*, the aetiologic agent of leprosy, but spill-over to other mammals such as nine-banded armadillos and red squirrels occurs. Although naturally acquired leprosy has also been described in captive nonhuman primates, the exact origins of infection remain unclear. Here, we report on leprosy-like lesions in two wild populations of western chimpanzees (*Pan troglodytes verus*) in the Cantanhez National Park, Guinea-Bissau, and the Taï National Park, Côte d’Ivoire, West Africa. Longitudinal monitoring of both populations revealed the progression of disease symptoms compatible with advanced leprosy. Screening of faecal and necropsy samples confirmed the presence of *M. leprae* as the causative agent at each site and phylogenomic comparisons with other strains from humans and other animals show that the chimpanzee strains belong to different and rare genotypes (4N/O and 2F). The independent evolutionary origin of *M. leprae* in two geographically distant populations of wild chimpanzees, with no prolonged direct contact with humans, suggests multiple introductions of *M. leprae* from an unknown animal or environmental source.

## MAIN

Leprosy is a neglected tropical disease caused by the bacterial pathogens *Mycobacterium leprae* and the more recently discovered *M. lepromatosis*^1,2^. In humans, the disease presents as a continuum of clinical manifestations with skin and nerve lesions of increasing severity, from the mildest tuberculoid form (or paucibacillary) to the most severe lepromatous type (or multibacillary)^3^. Symptoms develop after a long incubation period ranging from several months to 30 years, averaging five years in humans. As a result of sensory loss, leprosy can lead to permanent damage and severe deformity^4^. While leprosy prevalence has markedly decreased over the past decades, approximately 210,000 new human cases are still reported every year, of which 2.3% are located in West Africa^5^. Transmission is thought to occur primarily between individuals with prolonged and close contact via aerosolised nasal secretions and entry through nasal or respiratory mucosae, but the exact mechanism remains unclear^6,7^. The role of other routes, such as skin-to-skin contact, is unknown.

Leprosy-causing bacteria were once thought to be obligate human pathogens^8^. However, they can circulate in other animal hosts in the wild, such as in nine-banded armadillos (*Dasypus novemcinctus*) in the Americas and red squirrels (*Sciurus vulgaris*) in the United Kingdom^9,10^. Although initial infection was most likely incidental and of human origin, secondary animal hosts can subsequently represent a source of infection to humans^10–14^. In captivity, nonhuman primates, such as chimpanzees (*Pan troglodytes*)^15^, sooty mangabeys (*Cercocebus atys*)^16,17^ and cynomolgus macaques (*Macaca fascicularis*)^18^, developed leprosy spontaneously (i.e. not through laboratory experiments). However, it is unknown whether these species also contract leprosy in the wild.

Here, we report leprosy infections and their disease course in two wild populations of western chimpanzees (*Pan troglodytes verus*) in Cantanhez National Park (CNP), Guinea-Bissau, and in Taï National Park (TNP), Côte d’Ivoire, using a combination of camera trap and veterinary monitoring (Fig 1; Supplementary Information Note 1). From the analyses of faecal samples and post-mortem tissues, we identified *M. leprae* as the causative agent of the lesions observed and determined the phylogenetic placement of the respective strains based on their complete genome sequences.

**Fig. 1.**
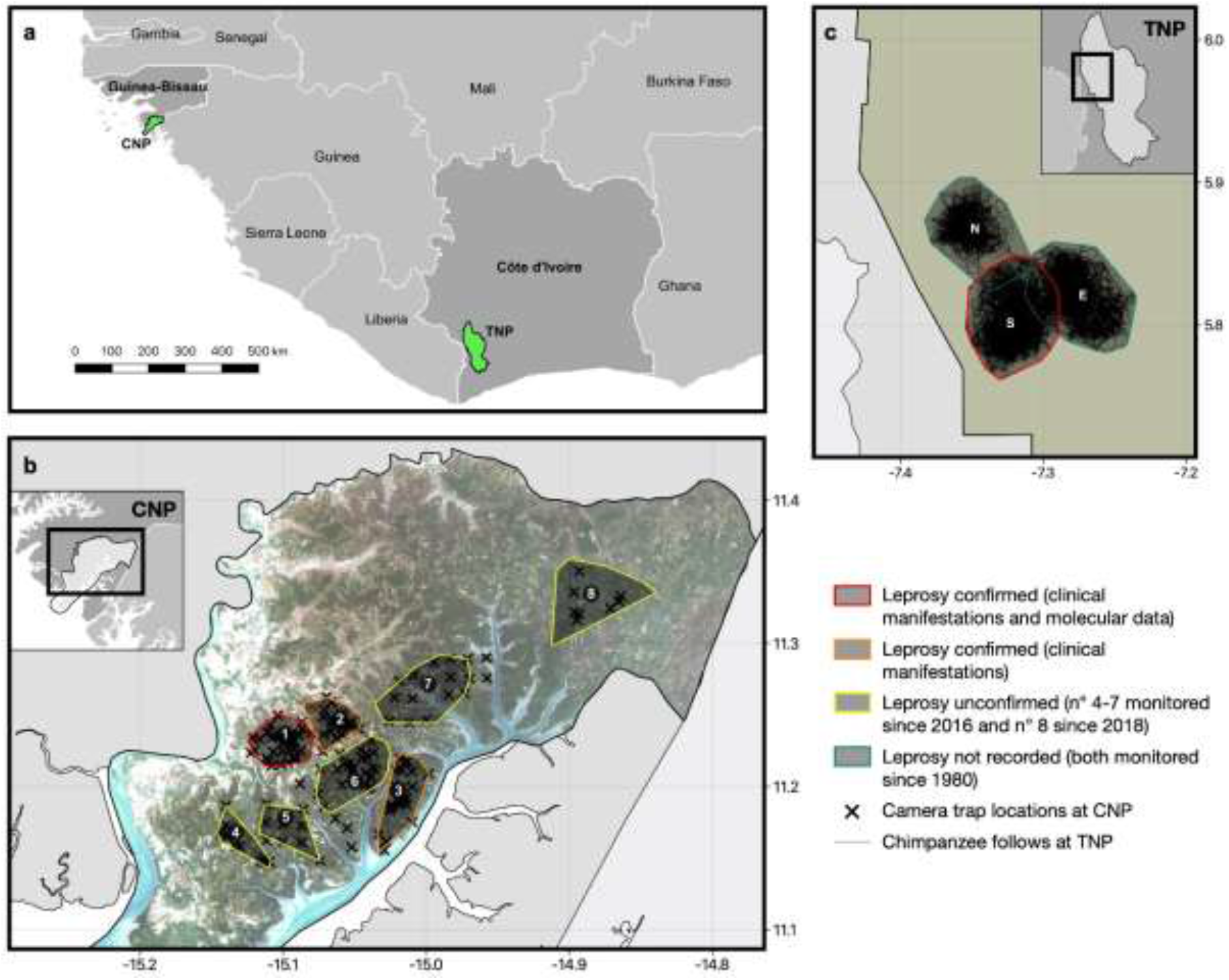
Maps of the chimpanzee study sites and chimpanzee communities. **a,** Map of the Cantanhez National Park (CNP), Guinea-Bissau, and the Taï National Park (TNP), Côte d’Ivoire, West Africa. **b,** Location of the chimpanzee communities at CNP that were monitored between 2015 and 2019 (1: Caiquene–Cadique; 2: Lautchande; 3: Cambeque; 4: Cabante; 5: Canamine; 6: Madina; 7: Amindara; 8: Guiledje). Estimated home ranges of chimpanzee communities at CNP are shown by 100% Minimum Convex Polygons of direct chimpanzee observations and indirect chimpanzee traces and nests during the study period. Red outline represents chimpanzee communities with at least one individual with clinical manifestations of leprosy, confirmed using molecular analysis; orange outline represents chimpanzee communities with at least one individual with clinical manifestations of leprosy; yellow colour represents monitored communities where clinical manifestations of leprosy have not been observed nor confirmed through molecular analysis. **c,** Location of the three habituated chimpanzee communities monitored at TNP (N: North; S: South; E: East). Estimated home ranges of chimpanzee communities at TNP are shown by 100% Minimum Convex Polygons of direct chimpanzee follows from December 2013 to October 2016. Red outline represents the community with individuals with clinical manifestations of leprosy, confirmed using molecular analysis and serological tests; blue colour represents communities where leprosy has not been recorded. CNP imagery is from Sentinel-2 (available at Sentinel Hub), and home range estimates were calculated in R using the package ‘adehabitatHR’^39^.

Chimpanzees at CNP are not habituated to human observers, precluding systematic behavioural observations. Longitudinal studies necessitate the use of camera traps, which we operated between 2015 and 2019. Of 624,194 data files (videos and photos) obtained across 211 locations at CNP (Fig 1b; Extended Data Table 1), 31,044 (5.0%) contained chimpanzees. The number of independent events (i.e. images separated by at least 60 minutes) totalled 4,336, and of these, 241 (5.6%) contained chimpanzees with severe leprosy-like lesions, including four clearly identifiable individuals (two adult females and two adult males) across three communities (Fig 1b; Extended Data Figs 1-2; Supplementary Information Note 2). As with humans, paucibacillary cases in chimpanzees may be present but easily go undetected. Since minor physical manifestations of leprosy are difficult to observe, they are not reported in our observations. All symptomatic chimpanzees showed hair loss and facial skin hypopigmentation, as well as plaques and nodules that covered different areas of their body (limbs, trunk and genitals), facial disfigurement and ulcerated and deformed hands (claw hand) and feet (Fig 2a-c), consistent with a multibacillary form of the disease. Longitudinal observations showed progression of symptoms across time with certain disease manifestations similar to those described in humans (e.g. progressive deformation of the hands for one individual) (Extended Data Fig 1; Supplementary Videos 1-3). To confirm infection with *M. leprae,* we collected faecal samples and tested them with two nested polymerase chain reaction (PCR) assays targeting the *M. leprae* specific RLEP repetitive element and 18kDa antigen gene. One out of 208 DNA-extracts from CNP was positive in both assays and a second was positive only in the more sensitive RLEP PCR^19^ (Table 1; Supplementary Table 1; Supplementary Information Note 3). Microsatellite analyses of the two positive samples confirmed that they originated from two distinct individuals, both of which were female (Supplementary Information Note 4; Extended Data Tables 2,3). Taken together, our results suggest that *M. leprae* is the most likely cause of a leprosy-like syndrome in chimpanzees from CNP.

**Fig. 2.**
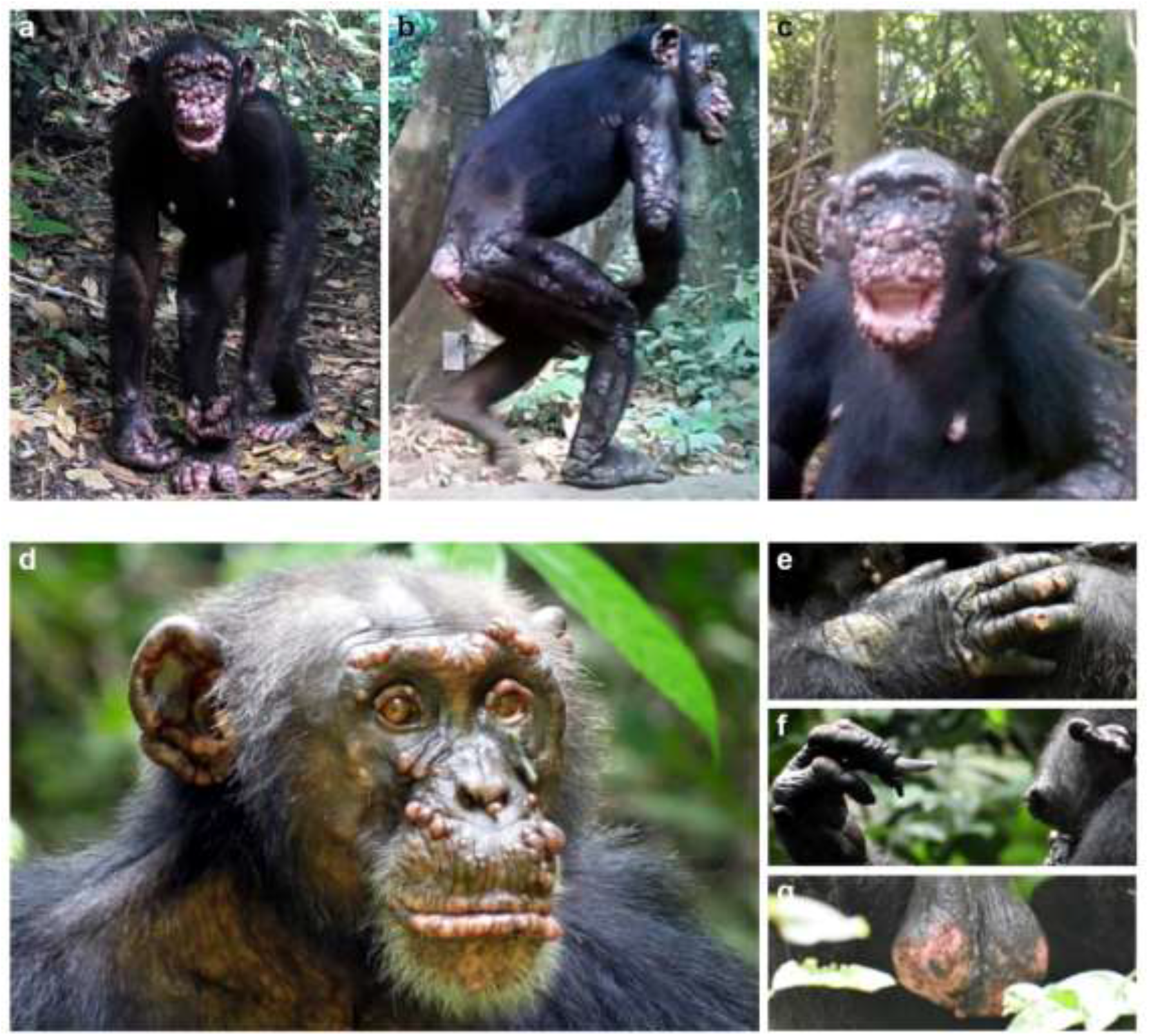
Clinical manifestations of leprosy in three chimpanzees at Cantanhez National Park (CNP), Guinea-Bissau, and the Taï National Park (TNP), Côte d’Ivoire. **a-c**, Clinical signs of leprosy in two adult female chimpanzees in CNP (images extracted from camera traps) **a**, Rita has large hypopigmented nodules covering the entire body; disfigurement of the face, ears, hand and feet (ulcerated lesions and swelling). **b**, Rita has extensive plaques covering all limbs, with hair loss. **c**, Brinkos has large hypopigmented nodules covering the entire face, with extreme disfigurement of the face and ears, and ulcerated plaques on the arms and the nipples. **d-g**, Clinical signs of leprosy in an adult male chimpanzee, Woodstock, at TNP **d**, Multiple hypopigmented nodules on the ears, brow ridges, eyelid margins, nostrils, lips and the area between the upper lip and the nose **e**, Hypopigmentation and swelling of the hands with ulcerations and hair loss on the dorsal side of the joints. **f**, Claw hand with nail loss and abnormal overgrowth of fingernails. **g**, Scrotal reddening and ulceration with fresh blood.

At TNP, chimpanzees are habituated to the presence of researchers and have been followed daily since 1979. In addition, systematic collection of necropsy samples has been performed on all dead individuals recovered in TNP since 2000. In June 2018, researchers first noticed leprosy-like lesions on Woodstock, an adult male chimpanzee from one of the three habituated communities (South) (Fig 1c; Extended Data Fig 3). Longitudinal observations through to 2020 showed that the initial small nodules seen on the ears, lips and under the eye became more prominent and were followed by several nodules on the eyebrows, eyelids, nostrils, ears, lips and face. Clinical manifestations later developed to include hypopigmentation of the skin on facial nodules, hands, feet and testicles, as well as the loss and abnormal growth of nails (Fig 2d-g). PCR screening of faecal samples collected from 2009 showed that *M. leprae* DNA was present in all samples collected since June 2018 (Table 1; Supplementary Table 1; Supplementary Information Notes 2 and 4). At this site, continuous non-invasive detection of *M. leprae* was associated with the onset and evolution of a leprosy-like disease.

Retrospective PCR screening of DNA extracted from all chimpanzee spleen samples (n=38 individuals) recovered from the collection of necropsy samples at TNP led to the identification of *M. leprae* DNA in two further individuals. An adult female from the same community named Zora, who had been killed by a leopard in 2009, tested positive in both PCR assays. The presence of *M. leprae* DNA was confirmed by PCR in various other organs of this chimpanzee (Table 1; Supplementary Table 1). Retrospective analyses of photos taken in the years before her death showed typical progressive skin hypopigmentation and nodule development since at least 2007 (Extended Data Fig 4). Formalin-fixed skin samples (hands and feet) were examined using hematoxylin and eosin, as well as Fite-Faraco, stains. In the histopathological examination, the skin presented typical signs of lepromatous leprosy characterised by a diffuse cutaneous cell infiltration in the dermis and the subcutis clearly separated from the basal layer of the epidermis (Extended Data Fig 5a). We detected moderate numbers of acid-fast bacilli, single or in clumps, within histiocytes, indicative of *M. leprae* (Extended Data Fig 5b). Since antibodies against the *M. leprae*-specific antigen phenolic glycolipid–I (PGL-I) are a hallmark of *M. leprae* infection in humans^20^, we also performed a PGL-I lateral flow rapid test^21^ on a blood sample from this individual, which showed strong seropositivity (Extended Data Fig 6). Faecal samples collected in the years before Zora’s death contained *M. leprae* DNA from 2002 onwards, implying at least seven years of infection (Table 1; Supplementary Information Table 1). In this case, disease manifestations, histopathological findings, serological and molecular data, as well as the overall course of the disease, all unambiguously point towards *M. leprae*-induced leprosy.

To ascertain whether other individuals in the South community of TNP were infected at the time of Zora’s death in 2009, cross-sectional screening of contact animals (n = 32) was performed by testing all available faecal samples (n = 176) collected in 2009 (Supplementary Information Table 1). Three other chimpanzees were PCR positive in single samples during this period, including Woodstock (Supplementary Information Note 4). Aside from Woodstock and Zora, clinical symptoms of leprosy have not been observed in any other individual at TNP despite extensive daily health and behavioural monitoring of all South community members over the last 20 years, and of neighbouring communities for the past 40 years^22,23^. Considering that, over this period, 467 individuals have been observed, it seems leprosy is a rare disease with low transmission levels in these chimpanzee communities.

To characterise the *M. leprae* strains causing leprosy in wild chimpanzees and to perform phylogenomic comparisons, we selected DNA extracts that were positive in both the RLEP and the less sensitive 18kDa PCR, which indicates relatively high levels of *M. leprae* DNA. For TNP, we only selected individuals that were positive in multiple samples. Following targeted enrichment using hybridization capture, samples were submitted to Illumina sequencing (Table 1; Supplementary Information Table 1). Sufficient coverage of *M*. *leprae* genomes was obtained for sample GB-CC064 (Guinea-Bissau) and for Zora (Côte d’Ivoire) with mean coverage of 39.3X and 25.8X, respectively (Table 1; Extended Data Table 4). We additionally generated 21 *M. leprae* genomes from human biopsies from five West African countries (Niger, Mali, Benin, Côte d’Ivoire and Senegal) and coverages ranged from 4.7X to 170X (Extended Data Table 4). We assembled a dataset, which included the genomes generated in this study and all previously available *M. leprae* genomes. Of the total 286 genomes, 64 originated from nine West African countries (Extended Data Fig. 7; Supplementary Information Note 5).

Bayesian and maximum-parsimony (MP) analyses (Fig 3; Supplementary Information Note 5) place the strain from Guinea-Bissau (GB-CC064) on branch 4, where it clusters outside the standard genotypes 4N, 4O and 4P, but within the so-called 4N/O genotype (Fig 3c)^24,25^. This 4N/O genotype is rare and comprises three *M. leprae* strains out of the 286 sequenced from worldwide. This includes only one strain (Ng13-33) from a patient in Niger (of 64 strains in West Africa), two strains (2188-2007 and 2188-2014) obtained from a single patient in Brazil (of 34 strains in Brazil)^26^, and two strains from two captive nonhuman primates originating from West Africa (CM1 and SM1)^25^ (Fig 3c). The branching order of these six strains was unresolved in our analyses, with a basal polytomy suggestive of star-like diversification within this genotype, and beyond within the group comprising all genotype 4 strains (4N/O, 4N, 4P and 4O) (Fig 3c). Divergence from the most recent common ancestor (MRCA) for this group is estimated to have occurred in the 6^th^ century C.E. (mean divergence time: 1437 years ago, 95% highest posterior density (HPD) 1132-1736 ya) (Fig 3a).

**Fig. 3.**
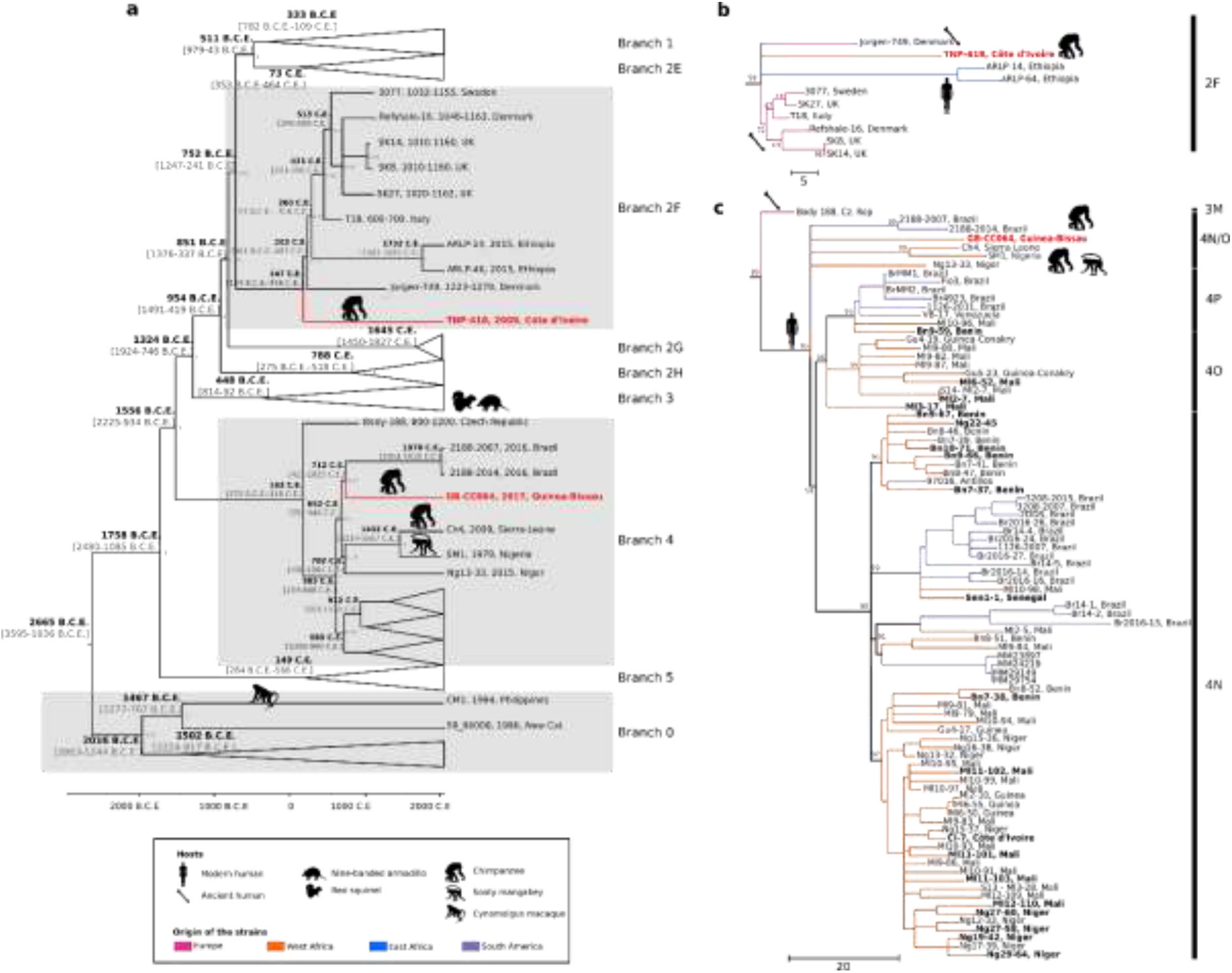
Phylogeny of *M. leprae* strains from human and animal hosts. **a**, Bayesian phylogenetic tree of 278 *M. leprae* genomes including the two new chimpanzee strains (in bold red). Hypermutated samples with mutations in the *nth* gene were excluded from the analysis. The tree is drawn to scale, with branch lengths representing years of age. Posterior probabilities are shown in grey. Some *M. leprae* branches are collapsed to increase readability. **b**, Maximum parsimony tree of the branch 2F. **c**, Maximum parsimony tree of the branch 4. The tree was initially constructed using 286 genomes (Supplementary Table 2), including the two new chimpanzee strains (in bold red) and 21 new genomes from West Africa (in bold), 500 bootstrap replicates and *M. lepromatosis* as outgroup. Sites with missing data were partially deleted (80% coverage cut-off), resulting in 4470 variable sites used for the tree calculation. Subtrees corresponding to branches were retrieved in MEGA7^40^. Corresponding genotypes are indicated on the side of each subtree. Samples are binned according to geographical origin as given in the legend. Animal silhouettes were available under Public Domain license at phylopic (http://phylopic.org/)

The strain that infected Zora in Côte d’Ivoire, designated TNP-418, belongs to branch 2F, within which branching order was also mostly unresolved (Fig 3a-b). The branch is currently composed of human strains from medieval Europe (n=7) and modern Ethiopia (n=2) (Fig. 3b), and this genotype has thus far never been reported in West Africa (Fig 3b). Bayesian analysis estimated a divergence time during the 2^nd^ century C.E. (mean of 1873 ya (95% HPD: 1564–2204 ya)), similar to previous predictions^27^. Samples of Woodstock did not yield sufficient genome coverage for phylogenomic analysis. However, SNPs recovered from Illumina reads and PCRs allowed us to assign this second *M. leprae* strain from Côte d’Ivoire to the same genotype as TNP-418 (Supplementary Information Note 6). Overall, phylogenomic analyses show that *M. leprae* strains in the chimpanzee populations of CNP and TNP are not closely related to one another.

The finding of *M. leprae*-induced leprosy in wild chimpanzee populations raises the question of the origin(s) of these infections. *M. leprae* is considered a human-adapted pathogen and earlier cases of leprosy affecting wildlife were compatible with anthroponosis. Therefore, the prime hypothesis would be a human-to-chimpanzee transmission. Potential routes of transmission include direct (e.g. skin-to-skin) contact and inhalation of respiratory droplets and/or fomites, with the assumption that, in all cases, prolonged and/or repeated exposure is required for transmission^4^. Chimpanzees at CNP are not habituated to human presence and are not approached at distances that would allow for transmission via respiratory droplets. Although chimpanzees at CNP inhabit an agroforest landscape and share access to natural and cultivated resources with humans^28^, present-day human-chimpanzee direct contact is uncommon. The exact nature of historic human-chimpanzee interactions at CNP remains, however, unknown. At TNP, direct human contact with wild chimpanzees has not been reported, with the South community distant from human settlements and agricultural areas. Human-to-animal transmission of pathogens has been shown at TNP^29,30^ but involved respiratory pathogens (pneumoviruses and human coronavirus OC43) that transmit easily and do not require prolonged exposure. In addition, *M. leprae* is thought to be transmitted from symptomatic humans^31^ and no leprosy cases have been reported among researchers or local research assistants. Combined with the rarity of the *M. leprae* genotypes detected in chimpanzees among human populations in West Africa, this suggests that recent human-to-chimpanzee transmission is unlikely.

The relatively old age of the lineages leading to the chimpanzee strains nevertheless raises the possibility of an ancient human-to-chimpanzee transmission. This hypothesis is, however, also unlikely for three reasons. First, the density of human populations at CNP and TNP 1,500-2,000 ya was even lower than it is currently, therefore reducing the likelihood of an ancient human-to-chimpanzee transmission. Second, two such human-to-chimpanzee transmission events would be required to explain our findings since *M. leprae* strains in CNP and TNP have distinct evolutionarily origins. Third, we can assume that if such transmission had occurred and the bacterium had persisted in chimpanzees, it should have spread more broadly as observed in *M. leprae*-infected squirrels and armadillos^10,12,13^. Therefore, ancient human-to-chimpanzee transmission is not a plausible mechanism to explain the presence of *M. leprae* in chimpanzee populations at CNP and TNP.

Alternatively, our findings may be explained by the presence of an unidentified animal or environmental source(s) that would serve as a leprosy reservoir. As chimpanzees hunt frequently, transmission may originate from their mammalian prey^32^. Nonhuman primates are the most hunted prey at TNP^33^, and possibly at CNP (Supplementary Information Note 3), but chimpanzees do occasionally consume other mammalian prey such as ungulates. If other animals were the proximal source of *M. leprae* infecting chimpanzees, the question of how they have become infected remains. An intriguing possibility is that an environmental source is at the origin of chimpanzee infections. Other mycobacteria can survive in water, including *M. ulcerans* and other non-tuberculous mycobacteria^34,35^, and molecular investigations have reported that *M. leprae* can survive in soil^36^. Experimental data also show that *M. leprae* multiplies in amoebae^37^, arthropods^38^, and ticks, which could contribute to the persistence of the bacteria in the environment. Taken together, our findings challenge the long-held assumption that humans are the main reservoir of *M. leprae* and suggest that this pathogen may sporadically emerge from environmental sources.

## Supporting information

Supplementary Table Titles

## METHODS

### Study sites

Observational study and sample collections were performed at the Cantanhez National Park (CNP) in southern Guinea-Bissau and the Taï National Park (TNP) in western Côte d’Ivoire (Fig 1a). CNP (1067 km^2^) comprises the Cubucaré peninsula in the sector of Bedanda, with the northeast of the park bordering the Republic of Guinea. The landscape at CNP consists of a mosaic of mainly mangroves, sub-humid forest patches, savannah grassland and woodland, remnant forest strips dominated by palm groves as well as agriculture. There are approximately 200 villages and settlements within the borders of the park, with an estimated human population of 24,000 individuals that comprise several ethnic groups^41^. Chimpanzees are not hunted for consumption within CNP due to local cultural beliefs and taboos^42^ but are sometimes killed in retaliation for foraging on crops^43,44^. There is a minimum of 12 chimpanzee communities at CNP^41^, all unhabituated to researchers, with approximately 35-60 individuals per community^45,46^. Numerous other wildlife taxa inhabit CNP, including six other nonhuman primate species^41,47^.

The TNP (5082 km^2^) consists of an evergreen lowland rainforest and is the largest remaining primary forest fragment in West Africa. It is home to a wide range of mammals that include 11 different nonhuman primate species^48,49^. There are no settlements or agricultural areas inside the National Park. As of October 2020, the three habituated communities, North, South and East, comprise 25, 39 and 35 individuals, respectively, although community sizes have varied over time. Systematic health monitoring of these communities has been ongoing since 2000^23^.

### Longitudinal observations and health monitoring

At CNP, camera traps (Bushnell Trophy Cam models 119774, 119877 and 119875) were deployed at 211 locations including across different habitat types (forest, mangrove-forest edge, orchards) within the home range of eight of the 12 putative chimpanzee communities (Fig 1b). Camera traps were set up over six data collection periods ranging from 2015 to 2019 (Extended Data Table 1). Targeted camera traps were deployed to record and monitor chimpanzee behaviour and disease occurrence. To maximise the chances of recording specific behaviours and identify leprosy-like symptoms in individuals, targeted camera traps were set up in locations that chimpanzees were known to use most often, sometimes in clusters, precluding uniform survey designs. Targeted camera traps were set up in video mode and active 24h per day. When triggered, targeted cameras recorded 10 to 60s of video with a minimum interval of 0.6s or 2s, depending on the camera trap model. Furthermore, systematically placed camera traps were used to obtain measures of wildlife occurrence and habitat use across the heterogeneous landscape^41^. Systematic camera traps were deployed across central CNP, at a minimum distance of 1km between sampling points, as well as within the home range of one chimpanzee community (Caiquene-Cadique) and were spaced at least 500m from one another. The camera traps pointed towards animal paths (often chimpanzee paths), small human paths also used by wildlife, and other areas presenting signs of animal activity. Systematic camera traps were set up to record three consecutive photographs when triggered. The Global Positioning System (GPS) coordinates, habitat type, date, time, and site description were recorded when setting up individual camera traps (targeted and systematic). Opportunistic observations of chimpanzees at CNP were made in 2013, during which chimpanzees were photographed and/or filmed using digital cameras.

Chimpanzees at TNP are fully habituated to human observers and all individuals in the habituated communities are individually identified. Behavioural and health monitoring of chimpanzees at TNP involves daily observation of habituated individuals by an interdisciplinary team comprising primatologists and veterinarians; investigations of wildlife mortality causes through necropsies on all animal carcasses found in the research area; and the collection of non-invasive samples such as faecal samples, laboratory investigations and the communication of the results to the park management for corrective and preventive measures^22^. Abnormalities in behaviour or clinical signs of disease are immediately reported and followed by detailed observation by the on-site veterinarian. In order to reduce the risk of transmission of human diseases to the chimpanzees, stringent hygiene measures have been put in place, including an initial five days quarantine for observers, keeping a distance of at least 7 meters, obligatory wearing of masks, with only healthy observers allowed to work in the forest^50,51^.

### Faecal and necropsy sample collection

At CNP, chimpanzee faecal samples were collected between July 2017 and December 2018. The date and putative chimpanzee community were recorded for each faecal sample. As defecation was rarely observed and to prevent the collection of redundant samples from the same individual, we avoided multiple samples found under the same chimpanzee nest and paid special attention if multiple samples were found in proximity on trails^45,52,53^. All samples were collected with the aid of a wooden spatula and stored at ambient temperature in 15ml tubes containing NAP buffer^54^. All samples were sent to the Robert Koch Institute for laboratory analysis. Even though chimpanzee faeces are easily distinguishable from those of other species, and were found in areas where chimpanzees had recently been present with associated signs such as feeding remains or knuckle prints, we genetically confirmed the presence of chimpanzee DNA in faecal samples that tested positive in either of the *M. leprae* PCRs or the mammal PCR for diet analysis (Supplementary Information Note 3).

At TNP, the long-term health monitoring program includes continuous collection of faecal and urine samples from known adult chimpanzees. Faeces are collected right after defecation, transferred in 2ml cryotubes with the aid of a plastic spatula and frozen in liquid nitrogen the same day. A full necropsy is systematically performed by the on-site veterinarian under high-level safety measures on all chimpanzees found dead. Necropsies follow a standardised biosafety protocol due to the occurrence of anthrax, Ebola and monkeypox in the area, that includes the use of full personal protective equipment and rigorous disinfection measures. Tissue samples of several internal organs are taken, as far as the state of carcass decomposition allows. After collection, all samples are first stored in liquid nitrogen and subsequently shipped on dry ice to the Robert Koch Institute for analyses.

### DNA extraction from faeces and necropsy samples

DNA extractions were performed at the Robert Koch Institute in a laboratory that has never been used for molecular *M. leprae* investigations. DNA was extracted from faecal and necropsy samples using the GeneMATRIX stool DNA purification kit (EURx, Poland) and the DNeasy Blood and Tissue kit (Qiagen, Germany), respectively, following the manufacturers’ instructions. Extracted DNA was then quantified using the Qubit™ dsDNA HS Assay kit (Thermo Fisher Scientific, MA, USA) and subsequently stored at −20°C until further use.

### Genetic identification of samples from infected chimpanzees at CNP

To determine whether faecal samples positive for *M. leprae* belonged to one or two individuals of CNP, we amplified chimpanzee DNA at 11 microsatellite loci and one sexing marker^55^. Due to the small quantity of starting DNA, not all loci were amplified and in some cases the amplification quality was low, impacting our ability to confidently interpret allele peak profiles (e.g. sample GB-CC064 failed to amplify for 5 out of the 11 loci) (Supplementary Information Note 4).

### Molecular screening of *M. leprae* in faecal and necropsy samples

*M. leprae* DNA was searched for using two nested PCR systems targeting the distinct but conservative repetitive element RLEP and the 18-kDa antigen gene as previously described (Extended Data Table 5). As several copies of RLEP are present in the *M. leprae* genome, this assay is considered to be more sensitive than 18 kDa, for which there is only a single copy. To prevent contamination at the laboratory at RKI and to enable us to identify if it occurs, we followed the following procedures: (1) separate rooms were used for preparation of PCR master mixes and the addition of DNA in the primary PCR; (2) the addition of the primary PCR product in the nested PCR, and (3) dUTPs were used for all PCRs instead of dNTPs. For both assays, primary PCRs were performed in 20μL reactions: up to 200ng of DNA was amplified using 1.25U of high-fidelity Platinum Taq™ polymerase (Thermo Fisher Scientific, MA, USA), 10x PCR buffer, 200μM dUTPs, 4mM MgCl2, and 200nM of both forward and reverse primers. The thermal cycling conditions for the primary and nested PCRs were as follows; denaturation at 95°C for 3 min, followed by 50 cycles of 95°C for 30 sec, 55°C (18kDa primers) or 58°C (RLEP primers) for 30 sec, and 72°C for 1 min as well as an elongation step at 72°C for 10 min. For the nested PCRs, 2μL of a 1:20 dilution of the primary PCR product was used as a template. Molecular grade water was used as a non-template control. PCR products were visualized on a 1.5% agarose gel stained with GelRed® (Biotium, CA, USA). Bands of the expected size were purified using the Purelink Gel extraction kit (Thermo Fisher Scientific, MA, USA). Both RLEP and 18-kDa nested PCR products are too short for direct Sanger sequencing. Therefore, fusion primers (primary PCR primers coupled with M13F and M13R primers) (Extended Data Table 5) were used for further amplification of the cleaned PCR products, applying the same conditions as in the primary PCR, but running only for 25 cycles. The resulting extended PCR products were then enzymatically cleaned using the ExoSAP-IT™ PCR Product Cleanup assay (Thermo Fisher, MA, USA) and Sanger sequenced using M13 primers. Resulting sequences were compared to publicly available nucleotide sequences using the Basic Local Alignment Search Tool (BLAST)^56^.

### Histopathology

To further confirm the infection, skin samples were sent to the German Primate Center in Göttingen, Germany for histopathological analyses. Samples were immersion-fixed in 10% neutral buffered formalin, embedded in paraffin, and stained with standard hematoxylin & eosin (HE) using the Varistain Gemini staining automat (Thermo Fisher Scientific, MA, USA). Samples were also stained with Fite-Faraco stain for the identification of acid-fast bacilli.

### Serology

A whole blood sample from Zora collected during the necropsy in 2009 was tested for the presence of the *M. leprae*-specific anti-Phenolic Glycolipid-I (PGL-I) antibodies using a chromatographic immunoassay developed for use with human blood following the instructions provided by the test manufacturers with a 1:10 diluted whole-blood sample. This rapid lateral flow test was produced by Dr. R. Cho using the synthetic ND-O-BSA antigen with financial support of the NIH/NIAID Leprosy Research Materials contract AI-55262 at Colorado State University. Test results were interpreted at five and 10 minutes. Human serum from a multibacillary leprosy patient donated by Prof. Spencer, Colorado State University, was used as a positive control.

### Library preparation, genome-wide capture and high-throughput sequencing for nonhuman primate samples (RKI)

Selected *M. leprae* positive faecal and necropsy samples (Supplementary Information Table 1) were converted into dual-indexed libraries using the NEBNext® Ultra™ II DNA Library Prep kit (New England Biolabs, MA, USA)^57,58^. To reconstruct whole genomes, libraries were target-enriched for *M. leprae* DNA using in-solution hybridisation capture with 80 nt RNA baits designed to cover the whole *M. leprae* genome (2-fold tiling; design can be shared upon request to the corresponding authors) and following the myBaits protocol as previously described^25^. Around 1.5μg of each DNA library was captured in single or pooled reactions. Two rounds of 24h hybridisation capture were performed followed by a post amplification step for each using the KAPA HiFi HotStart Library amplification kit with 12 to 16 cycles to generate around 200ng of enriched library per sample. Finally, enriched libraries were purified using the silica based MinElute reaction cleanup kit (Qiagen, Germany) followed by quantification with the KAPA library quantification kit (Roche, Switzerland). Libraries were then normalized and pooled across sequencing lanes on an Illumina NextSeq 500 mid output kit v2; 300 cycles (Illumina, CA, USA).

### Human specimens: sample collection, DNA extraction, library preparation, genome-wide capture and high-throughput sequencing (EPFL)

Samples (skin biopsies or DNA extracts) from leprosy patients from five West African countries with positive bacillary index, Niger (n=5), Mali (n=8), Benin (n=6), Côte d’Ivoire (n=1) and Senegal (n=1), were obtained from the respective National Leprosy Control Programmes in the framework of the leprosy drug resistance surveillance programs or from previous investigation^59^.

DNA was extracted from skin biopsies using the total DNA extraction method as described previously^60^. DNA was quantified with a Qubit fluorometer using the Qubit™ dsDNA BR Assay kit (Thermo Fisher Scientific, MA, USA) prior library preparation. DNA libraries were prepared using the Kapa Hyper Prep kit (Roche, Switzerland) as per the manufacturer’s recommendation using Kapa Dual Indexed Adapter (Roche, Switzerland) followed by in-solution capture enrichment with 80nt RNA baits with 2x tiling density for 48h at 65°C as described recently^60^. Post-capture amplification was performed with seven cycles. Enriched libraries were purified using a 1X ratio of KAPA Pure beads (Roche, Switzerland) followed by quantification with the KAPA library quantification kit (Roche, Switzerland) and quality control of the fragment with the Agilent 2200 TapeStation (Agilent Technologies, CA, USA). Libraries were then normalized and pooled across sequencing lanes on an Illumina NextSeq 500 on a high output kit v2; 75 cycles (Illumina, CA, USA).

### Genomic data analysis

Raw reads were processed as described elsewhere^24^. Putative unique variants of GB-CC064 and TNP-418 strains were manually checked and visualized using the Integrative Genomics Viewer^61^.

### Genome-wide comparison and phylogenetic tree

SNPs of the two newly sequenced genomes from chimpanzees were compared to the 263 publicly available *M. leprae* genomes (Supplementary Table 2)^25,60,62–64^ and 21 new genomes from West African countries (Supplementary Information Note 5). Phylogenetic analyses were performed using a concatenated SNP alignment (Supplementary Table 3). Maximum Parsimony (MP) trees were constructed in MEGA7^40^ with the 286 genomes available (Supplementary Table 2) using 500 bootstrap replicates and *M. lepromatosis*^65^ as outgroup. Sites with missing data were partially deleted (80% coverage cut-off), resulting in 4470 variable sites used for the tree calculation.

### Dating analysis

Dating analyses were done using BEAST2 (v2.5.2)^66^ as described previously with 278 genomes and an increased chain length from 50 to 100 million^24^. Briefly, the concatenated SNPs for each sample were used for tip dating analysis (Supplementary Information Table 4). Hypermutated strains and highly mutated genes associated with drug resistance (in yellow, Supplementary Information Table 3) were omitted^24,60^, manual curation of the MP and BEAST input file was done at the positions described in Supplementary Information Table 5 for GB-CC064 and TNP-418. Sites with missing data as well as constant sites were included in the analysis, as previously described^24^. Only unambiguous constant sites, i.e., loci where the reference base was called in all samples, were included.

### PCR genotyping of insufficiently covered *M. leprae* genomes from positive chimpanzees

The genome coverage for the strain infecting Woodstock was low. To be able to determine the genotype, we identified specific variants from the genome-wide comparison of TNP-418 (the strain infecting Zora, an individual from the same social group) with other strains from branch 2F (Supplementary Table 6). Variants were manually checked and visualized in the partially covered genome from the strain infecting Woodstock using IGV software (Supplementary Table 6). Two variants not covered by high throughput sequencing data were also selected for specific PCR screening. Primers were designed using the Primer3 web tool (http://bioinfo.ut.ee/primer3-0.4.0/) based on the Mycobrowser sequences^67^ and are described in Extended Data Table 5. All PCR conditions were the same as in the *M. leprae* screening PCRs except for the primer sets and associated annealing temperatures (Extended Data Table 5).

## ACKNOWLEDGEMENTS

We thank the Instituto da Biodiversidade e das Areas Protegidas (IBAP) for their permission to conduct research in Guinea-Bissau and for logistical support. A special thank goes to Queba Quecuta, Director of CNP, and research assistants and local guards, in particular Mamadu Cassamá, Iaia Tawél Camará, Djibi Indjai, Serifu Sila, Alia Camará, Idrissa Galiza, Fernando Ndafa, Adulai Camará, Braima Sedja Vieira for assisting with data collection and providing invaluable advice in Cantanhez National Park. We thank village chiefs, Nalu leaders and Régulos for granting us permission to conduct research.

We thank the Ministère de l’Enseignement Supérieur et de la Recherche Scientifique and the Ministère des Eaux et Forêts in Côte d’Ivoire, and the Office Ivoirien des Parcs et Réserves for permitting the research at TNP. We are grateful to the staff of the Taï Chimpanzee Project and the Centre Suisse de Recherches Scientifiques for their constant support of our work in TNP. Thanks to Sylvain Lemoine for providing shapefiles on chimpanzee home ranges at Taï National Park.

We would like to thank Prof. John McKinney, Dr Neeraj Dhar, Suzanne Balharry, Dr. Bastien Mangeat and the Gene expression Core Facility from the Ecole Polytechnique Fédérale de Lausanne for support. The authors would like to thank Prof. Anne Stone from Arizona State University for her constructive comments on the manuscript. Finally, the authors are grateful to all the patients and clinical staff who participated in the study.

## Data Availability Statement

Sequence data are available from the NCBI Sequence Read Archive (SRA) Bioproject PRJNA664360 Biosamples SAM16207289-16207321. Biosample codes for all samples used in this study are given in the Supplementary Data. Other relevant data supporting the findings of the study are available in this published article and its Supplementary Information files, or from the corresponding author upon request.

## Author contributions

Collection and analysis of chimpanzee camera trap data (CNP): KH, EB, JB, MR

Collection of long-term data on chimpanzees and their health (TNP): RMW, FHL

Collection of chimpanzee samples (CNP and TNP): KH, EB, JB, MR, KP, AD, SM, FHL

Logistics and fieldwork: KH, AR, AS, EC-H, RMW, FHL

Performance of necropsies: SM, AD, FHL

Collection and provision of human samples: MG, RCJ, MK, SOS, STC, HZ

DNA extraction, library preparation, enrichment and whole genome sequencing: BM, CA, PB

PCR and SNP confirmation: BM, LVP, CA

Chimpanzee microsatellite analysis: CB, JL-J

Computational analysis: CA, SCS, BM

Dating analysis: AB

Serological Investigations: BM

Provide material and protocol for serological investigation: JSS

Funding acquisition for leprosy research: KH, CA, JSS, CB, STC, RMW, FHL

KH, CA, CB, SCS, FL wrote the manuscript with considerable input from BM, KP, EB, JB, MR, AD, LVP, with contributions from all authors.

All authors approved the submitted manuscript.

The authors declare no competing interests.

## Additional information

Supplementary Information is available for this paper.

Correspondence and requests for materials should be addressed to FHL

**Extended data (see corresponding file)**

## References main text

1. Han, X. Y. et al. A new *Mycobacterium* species causing diffuse lepromatous leprosy. Am. J. Clin. Pathol. 130, 856–864 (2008).

2. Han, X. Y. et al. Comparative sequence analysis of *Mycobacterium leprae* and the new leprosy-causing *Mycobacterium lepromatosis*. J. Bacteriol. 191, 6067–6074 (2009).

3. Ridley, D. S. & Jopling, W. H. Classification of leprosy according to immunity. A five-group system. Int. J. Lepr. Mycobact. Dis. Off. Organ Int. Lepr. Assoc. 34, 255–273 (1966).

4. Britton, W. J. & Lockwood, D. N. Leprosy. The Lancet 363, 1209–1219 (2004).

5. WHO. Leprosy (Hansen’s disease). https://www.who.int/westernpacific/health-topics/leprosy (2020).

6. Araujo, S., Freitas, L. O., Goulart, L. R. & Goulart, I. M. B. Molecular evidence for the aerial route of infection of *Mycobacterium leprae* and the role of asymptomatic carriers in the persistence of leprosy. Clin. Infect. Dis. Off. Publ. Infect. Dis. Soc. Am. 63, 1412–1420 (2016).

7. Lastória, J. C. & Abreu, M. A. M. M. de. Leprosy: review of the epidemiological, clinical, and etiopathogenic aspects - Part 1. An. Bras. Dermatol. 89, 205–218 (2014).

8. Walker, S. L., Withington, S. G. & Lockwood, D. N. J. 41 - Leprosy. in Manson’s tropical infectious diseases (twenty-third Edition) (eds. Farrar, J. et al.) 506–518.e1 (W.B. Saunders, 2014). doi:10.1016/B978-0-7020-5101-2.00042-X.

9. Truman, R. Leprosy in wild armadillos. Lepr. Rev. 76, 198–208 (2005).

10. Avanzi, C. et al. Red squirrels in the British Isles are infected with leprosy bacilli. Science 354, 744–747 (2016).

11. Monot, M. et al. On the origin of leprosy. Science 308, 1040–1042 (2005).

12. Truman, R. W. et al. Probable zoonotic leprosy in the southern United States. N. Engl. J. Med. 364, 1626–1633 (2011).

13. Sharma, R. et al. Zoonotic leprosy in the southeastern United States. Emerg. Infect. Dis. 21, 2127–2134 (2015).

14. Silva, M. B. da et al. Evidence of zoonotic leprosy in Pará, Brazilian Amazon, and risks associated with human contact or consumption of armadillos. PLoS Negl. Trop. Dis. 12, e0006532 (2018).

15. Suzuki, K., Tanigawa, K., Kawashima, A., Miyamura, T. & Ishii, N. Chimpanzees used for medical research shed light on the pathoetiology of leprosy. Future Microbiol. 6, 1151–1157 (2011).

16. Meyers, W. M. et al. Leprosy in a mangabey monkey--naturally acquired infection. Int. J. Lepr. Mycobact. Dis. Off. Organ Int. Lepr. Assoc. 53, 1–14 (1985).

17. Gormus, B. J. et al. A second sooty mangabey monkey with naturally acquired leprosy: first reported possible monkey-to-monkey transmission. Int. J. Lepr. Mycobact. Dis. Off. Organ Int. Lepr. Assoc. 56, 61–65 (1988).

18. Valverde, C. R., Canfield, D., Tarara, R., Esteves, M. I. & Gormus, B. J. Spontaneous leprosy in a wild-caught cynomolgus macaque. Int. J. Lepr. Mycobact. Dis. Off. Organ Int. Lepr. Assoc. 66, 140–148 (1998).

19. Donoghue, H. D., Holton, J. & Spigelman, M. PCR primers that can detect low levels of *Mycobacterium leprae* DNA. J. Med. Microbiol. 50, 177–182 (2001).

20. Stefani, M. M. et al. Assessment of anti-PGL-I as a prognostic marker of leprosy reaction. Int. J. Lepr. Mycobact. Dis. Off. Organ Int. Lepr. Assoc. 66, 356–364 (1998).

21. Spencer, J. S. & Brennan, P. J. The role of *Mycobacterium leprae* phenolic glycolipid I (PGL-I) in serodiagnosis and in the pathogenesis of leprosy. Lepr. Rev. 82, 344–357 (2011).

22. Leendertz, F. H. et al. Pathogens as drivers of population declines: the importance of systematic monitoring in great apes and other threatened mammals. Biol. Conserv. 131, 325–337 (2006).

23. The chimpanzees of the Taï Forest: 40 years of research. (Cambridge University Press, 2019). doi:10.1017/9781108674218.

24. Benjak, A. et al. Phylogenomics and antimicrobial resistance of the leprosy bacillus *Mycobacterium leprae*. Nat. Commun. 9, 352 (2018).

25. Honap, T. P. et al. *Mycobacterium leprae* genomes from naturally infected nonhuman primates. PLoS Negl. Trop. Dis. 12, (2018).

26. Stefani, M. M. A. et al. Whole genome sequencing distinguishes between relapse and reinfection in recurrent leprosy cases. PLoS Negl. Trop. Dis. 11, e0005598 (2017).

27. Schuenemann, V. J. et al. Ancient genomes reveal a high diversity of *Mycobacterium leprae* in medieval Europe. PLoS Pathog. 14, e1006997 (2018).

28. Hockings, K. J., Parathian, H., Bessa, J. & Frazão-Moreira, A. Extensive overlap in the selection of wild fruits by chimpanzees and humans: implications for the management of complex social-ecological systems. Front. Ecol. Evol. 8, (2020).

29. Köndgen, S. et al. Pandemic human viruses cause decline of endangered great apes. Curr. Biol. 18, 260–264 (2008).

30. Patrono, L. V. et al. Human coronavirus OC43 outbreak in wild chimpanzees, Côte d’Ivoire, 2016. Emerg. Microbes Infect. 7, (2018).

31. Richardus, J. H., Ignotti, E. & Smith, W. C. S. Epidemiology of leprosy, chapter 1.1. in International Textbook of Leprosy (eds. Scollard, D. M. & Gillis, T. P.) (2016).

32. Gogarten, J. F. et al. The ecology of primate retroviruses – an assessment of 12 years of retroviral studies in the Taï national park area, Côte d’Ivoire. Virology 0, 147–153 (2014).

33. Boesch, C. & Boesch, H. Hunting behavior of wild chimpanzees in the Taï National Park. Am. J. Phys. Anthropol. 78, 547–573 (1989).

34. Stinear, T. et al. Identification of *Mycobacterium ulcerans* in the environment from regions in Southeast Australia in which it is endemic with sequence capture-PCR. Appl. Environ. Microbiol. 66, 3206–3213 (2000).

35. Rabinowitz, P. & Conti, L. Human-animal medicine: clinical approaches to zoonoses, toxicants and other shared health risks. (Elsevier, 2010).

36. Lahiri, R. & Adams, L. B. Cultivation and viability determination of *Mycobacterium leprae*. in International Textbook of Leprosy (2016).

37. Wheat, W. H. et al. Long-term survival and virulence of *Mycobacterium leprae* in amoebal cysts. PLoS Negl. Trop. Dis. 8, e3405 (2014).

38. Neumann, A. da S. et al. Experimental infection of *Rhodnius prolixus* (Hemiptera, Triatominae) with *Mycobacterium leprae* indicates potential for leprosy transmission. PLoS ONE 11, e0156037 (2016).

39. Calenge, C. The package adehabitat for the R software: a tool for the analysis of space and habitat use by animals. Ecol. Model. 197, 516–519 (2006).

40. Kumar, S., Stecher, G. & Tamura, K. MEGA7: molecular evolutionary genetics analysis version 7.0 for bigger datasets. Mol. Biol. Evol. 33, 1870–1874 (2016).

## References methods

41. Bersacola, E. Zooming in on human-wildlife coexistence: primate community responses in a shared agroforest landscape in Guinea-Bissau. (Oxford Brookes University, 2019).

42. Parathian, H. E., McLennan, M. R., Hill, C. M., Frazão-Moreira, A. & Hockings, K. J. Breaking through disciplinary barriers: human–wildlife interactions and multispecies ethnography. Int. J. Primatol. (2018) doi:10.1007/s10764-018-0027-9.

43. Sousa, C. M. A. M. R. & Frazão-Moreira, A. Etnoprimatologia ao serviço da conservação na Guiné-Bissau: o chimpanzé como exemplo. in Etnoecologia em perspectiva: natureza, cultura e conservação (eds. Alves, A. G. C., Souto, F. J. B. & Peroni, N.) 187–200 (NUPEEA, 2010).

44. Sousa, J., Vicente, L., Gippoliti, S., Casanova, C. & Sousa, C. Local knowledge and perceptions of chimpanzees in Cantanhez National Park, Guinea-Bissau. Am. J. Primatol. 76, 122–134 (2014).

45. Bessa, J., Sousa, C. & Hockings, K. J. Feeding ecology of chimpanzees (*Pan troglodytes verus*) inhabiting a forest-mangrove-savanna-agricultural matrix at Caiquene-Cadique, Cantanhez National Park, Guinea-Bissau. Am. J. Primatol. 77, 651–665 (2015).

46. Vieira, W. F., Kerry, C. & Hockings, K. J. A comparison of methods to determine chimpanzee home-range size in a forest–farm mosaic at Madina in Cantanhez National Park, Guinea-Bissau. Primates 60, 355–365 (2019).

47. Hockings, K. J. & Sousa, C. Human-chimpanzee sympatry and interactions in Cantanhez National Park, Guinea-Bissau: current research and future directions. Primate Conserv. 26, 57–65 (2013).

48. Monkeys of the Taï forest: an African primate community. (Cambridge University Press, 2007). doi:10.1017/CBO9780511542121.

49. Boesch, C. & Boesch-Achermann, H. The chimpanzees of the Taï Forest: behavioural ecology and evolution. (Oxford University Press, 2000).

50. Gilardi, K. V. K. et al. Best practice guidelines for health monitoring and disease control in great ape populations. (IUCN, 2015).

51. Grützmacher, K. et al. Human quarantine: toward reducing infectious pressure on chimpanzees at the Taï Chimpanzee Project, Côte d’Ivoire. Am. J. Primatol. 80, (2018).

52. McLennan, M. R. Diet and feeding ecology of chimpanzees (*Pan troglodytes*) in Bulindi, Uganda: foraging strategies at the forest–farm interface. Int. J. Primatol. 34, 585–614 (2013).

53. McCarthy, M. S. et al. Genetic censusing identifies an unexpectedly sizeable population of an endangered large mammal in a fragmented forest landscape. BMC Ecol. 15, 21 (2015).

54. Camacho-Sanchez, M., Burraco, P., Gomez-Mestre, I. & Leonard, J. A. Preservation of RNA and DNA from mammal samples under field conditions. Mol. Ecol. Resour. 13, 663–673 (2013).

55. Borges, F. F. A country-level genetic survey of the IUCN Critically Endangered western chimpanzee (*Pan troglodytes verus*) in Guinea-Bissau. (University of Porto, 2017).

56. Altschul, S. F., Gish, W., Miller, W., Myers, E. W. & Lipman, D. J. Basic local alignment search tool. J. Mol. Biol. 215, 403–410 (1990).

57. Mubemba, B. et al. Yaws disease caused by *Treponema pallidum* subspecies *pertenue* in wild chimpanzee, Guinea, 2019. Infect. Dis. J. - CDC 26, 1283–1286 (2020).

58. Patrono, L. V. et al. Monkeypox virus emergence in wild chimpanzees reveals distinct clinical outcomes and viral diversity. Nat. Microbiol. 5, 955–965 (2020).

59. Monot, M. et al. Comparative genomic and phylogeographic analysis of *Mycobacterium leprae*. Nat. Genet. 41, 1282–1289 (2009).

60. Avanzi, C. et al. Population genomics of *Mycobacterium leprae* reveals a new genotype in Madagascar and the Comoros. Front. Microbiol. 11, (2020).

61. Robinson, J. T., Thorvaldsdóttir, H., Wenger, A. M., Zehir, A. & Mesirov, J. P. Variant review with the integrative genomics viewer. Cancer Res. 77, e31–e34 (2017).

62. Avanzi, C. et al. Emergence of *Mycobacterium leprae* rifampin resistance evaluated by whole-genome sequencing after 48 years of irregular treatment. Antimicrob. Agents Chemother. 64, (2020).

63. Guan, Q., Almutairi, T. S., Alhalouli, T., Pain, A. & Alasmari, F. Metagenomics of imported multidrug-resistant *Mycobacterium leprae*, Saudi Arabia, 2017. Emerg. Infect. Dis. J. - CDC 26, 615–617 (2020).

64. Tió-Coma, M. et al. Genomic characterization of *Mycobacterium leprae* to explore transmission patterns identifies new subtype in Bangladesh. Front. Microbiol. 11, (2020).

65. Singh, P. et al. Insight into the evolution and origin of leprosy bacilli from the genome sequence of *Mycobacterium lepromatosis*. Proc. Natl. Acad. Sci. U. S. A. 112, 4459–4464 (2015).

66. Volz, E. M. & Siveroni, I. Bayesian phylodynamic inference with complex models. PLoS Comput. Biol. 14, e1006546 (2018).

67. Kapopoulou, A., Lew, J. M. & Cole, S. T. The MycoBrowser portal: a comprehensive and manually annotated resource for mycobacterial genomes. Tuberc. Edinb. Scotl. 91, 8–13 (2011).

