## Extended Data Figures and Tables for "Leprosy in wild chimpanzees"

### Extended data Figures & Tables

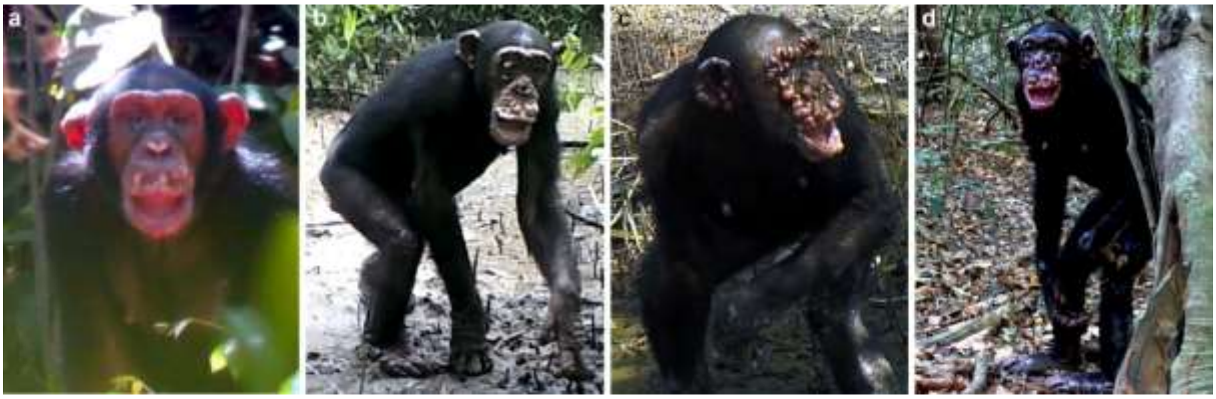


**Extended Data Fig. 1. Disease progression of leprosy in an adult female chimpanzee at CNP (Rita) over the course of 5 years. a**, 2013/05 – Hypopigmentation of skin around the mouth and nose, small nodule on the lower lip and left ear (opportunistically recorded with a video camera before the start of longitudinal health monitoring with camera traps). **b**, 2015/12 – Large nodules between the upper lip and nose, with multiple small nodules on the eyelids, cheek, ears margins, lower lip, and brow ridge. Small dry patches with hair loss on the wrists, knees and elbows. **c**, 2017/12 – Nodules increase in number, with apparent swelling and reddening, facial disfigurement, and claw hand. Plaques appear on the wrist, knee, and elbow joints, with an increase in hair thinning. **d**, 2018/05 – Face and ears completely covered by large nodules, with facial disfigurement and generalised hair loss on limbs and lower back. Nodule formation and swelling of fingers and toes, with disfigurement of hands and feet, and more severe claw hand. Some plaques on the body are ulcerated, and the individual has clear weight loss.

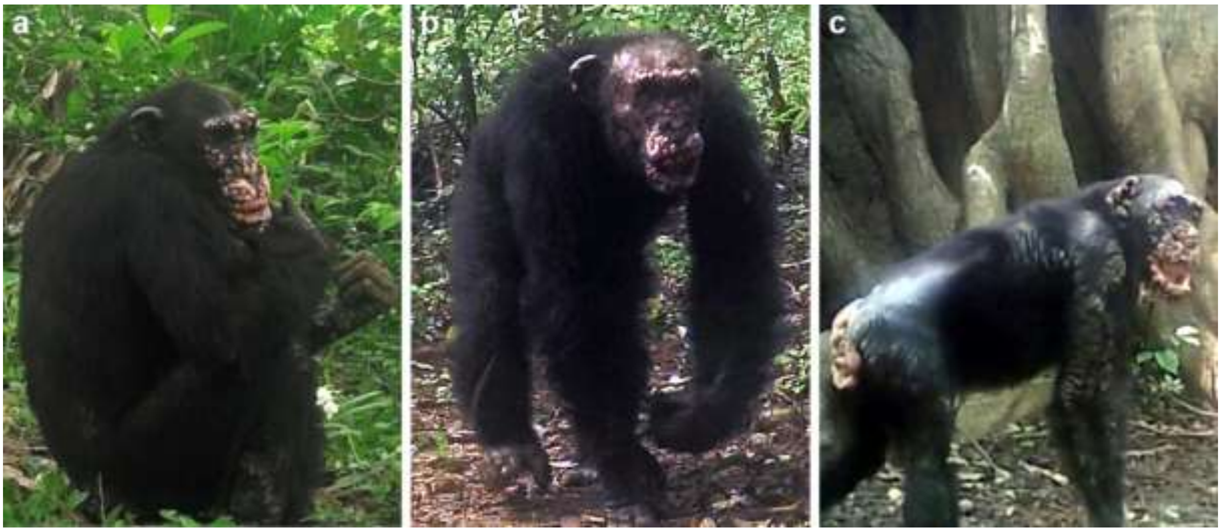


**Extended Data Fig. 2. Disease manifestations in three adult chimpanzees at CNP. a,** Jimi (Lautchande in 2018/06) – First observation of lesions in 2015. The head is completely covered with multiple nodules of reddish colour, some of which are ulcerated. Ear margins are thickened. Hands and feet present nodules and plaques, and the scrotum is affected (not visible on picture). **b,** Baaba (Cambeque in 2017/08) – First observation of lesions in 2017. Multiple hypopigmented nodules on the brow ridge, cheek, and upper and lower lips. Ears have thickened margins and nodules. There is hair thinning, with multiple small plaques present on the upper and lower limbs, back, abdomen and shoulders. **c,** Brinkos (Caiquene-Cadique in 2018/10) – First observation of lesions in 2015. Facial disfigurement, with the ulceration of nodules and a hanging lower lip. Hands and feet are ulcerated, and fingers are swollen. There are nodules on the nipples, and plaques covering the lower back, shoulders and arm are ulcerated, with hair loss.

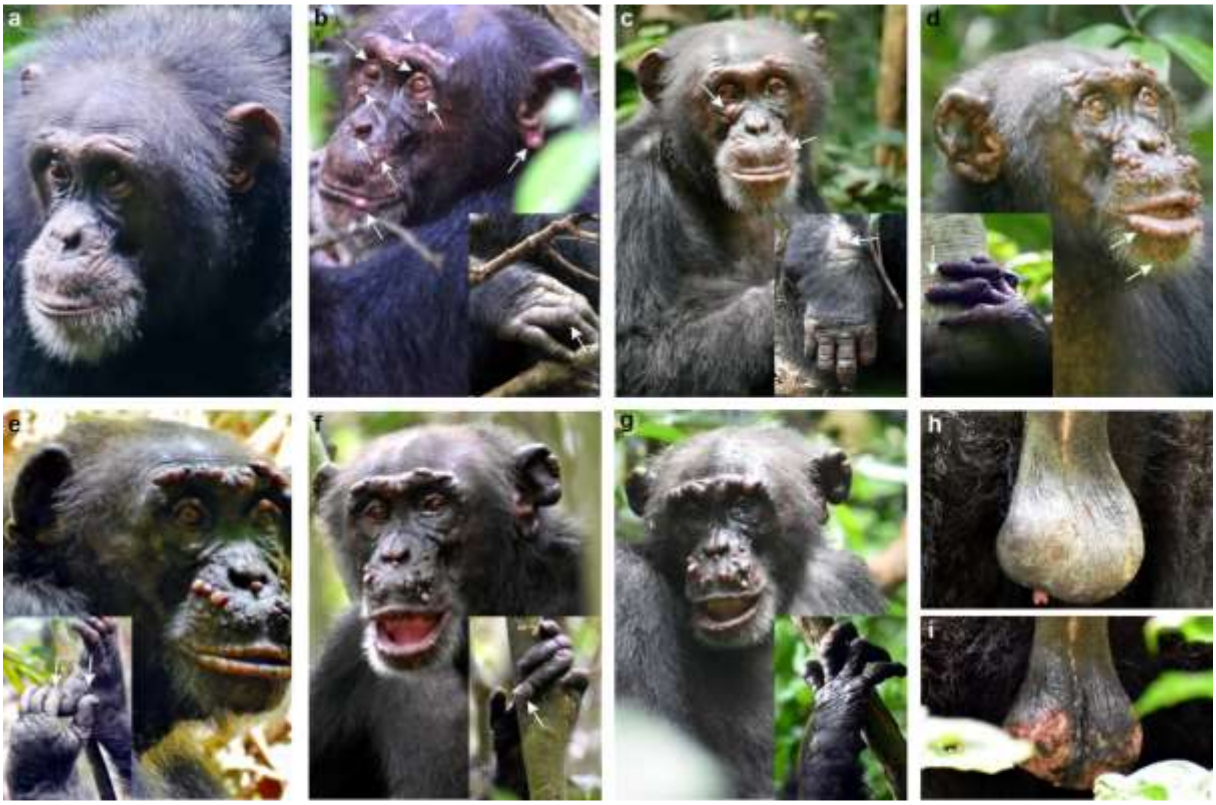

**Extended Data Fig. 3. Disease progression of leprosy in an adult male chimpanzee, Woodstock, at TNP, over the course of two years (2018-2020). a**, 2017/01 – Woodstock prior to the appearance of clinical signs. **b**, 2018/06 – First hypopigmented nodules appear on the face (see arrows), with swelling and hypopigmentation on both hands, and ulceration on the right hand. **c**, 2018/10 – Existing nodules increase in size and new smaller ones appear (see arrows). Development of mucopurulent discharge from the left eye, and lower eyelid is turned outward. Hair loss and ulceration on dorsal part of right wrist and hand. **d**, 2019/04 – Most existing nodules increase in size and become pedunculated, and the nodule under the eye shrinks, and several new nodules appear (see arrows). Suspected start of nasal involvement, and right ear starts to become disfigured. Both hands are slightly swollen and hypopigmented, with the loss of nail plate on the 4$^{th}$ finger of the left hand, and the 3$^{rd}$ and 5$^{th}$ fingers show early stage of abnormal nail overgrowth. **e,** 2019/10 – Facial lesions increase in size, and some become darkly pigmented. New lesions appear on the brow ridge, with nodules above the lips and between the lips, and the nose becomes pedunculated. The loss of nail plate, and nail bed becomes exposed on the 1$^{st}$ and 2$^{nd}$ fingers of the left hand. **f**, 2020/04 – In general, facial nodules appear smaller than before, and the nodule under the left eye disappears. On the left hand, the nail of the 4$^{th}$ finger shows an advanced stage of abnormal nail overgrowth, and the 3$^{rd}$ and 5$^{th}$ fingernails show early stage of abnormal nail overgrowth. **g**, 2020/07 – Facial nodules appear larger with many hypopigmented, and both ears are swollen and disfigured. Nasal involvement becomes apparent. Both hands are swollen and hypopigmented. Skin ulcerations present on the right hand, with possible claw hand on the left hand. **h**, 2019/04 – Slight hypopigmentation of scrotum. **i**, 2020/07 – Reddening and ulceration of scrotum; fresh blood observed.

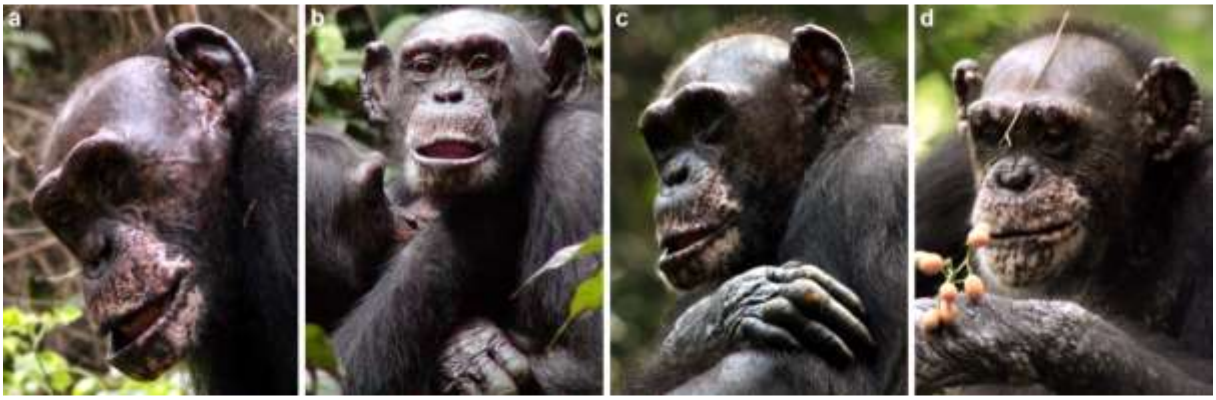


**Extended Data Fig. 4. Disease progression of leprosy in an adult female chimpanzee of TNP Zora over the course of two years. a,** 2007/12 – Zora prior to the appearance of clinical signs of leprosy. **b,** 2008/01 – Appearance of nodules on the right ear and both eyebrow ridges. **c,** 2008/12 – Appearance of nodules on the left ear, and ulceration of the skin at the 2$^{nd}$, 3$^{rd}$ and 4$^{th}$ proximal interphalangeal joint level of the right hand. **d,** 2009/04 – Nodular lesions on both ears and brow ridge appear aggravated, with nodular lesions on the lips, and above the mouth (four months prior to death).

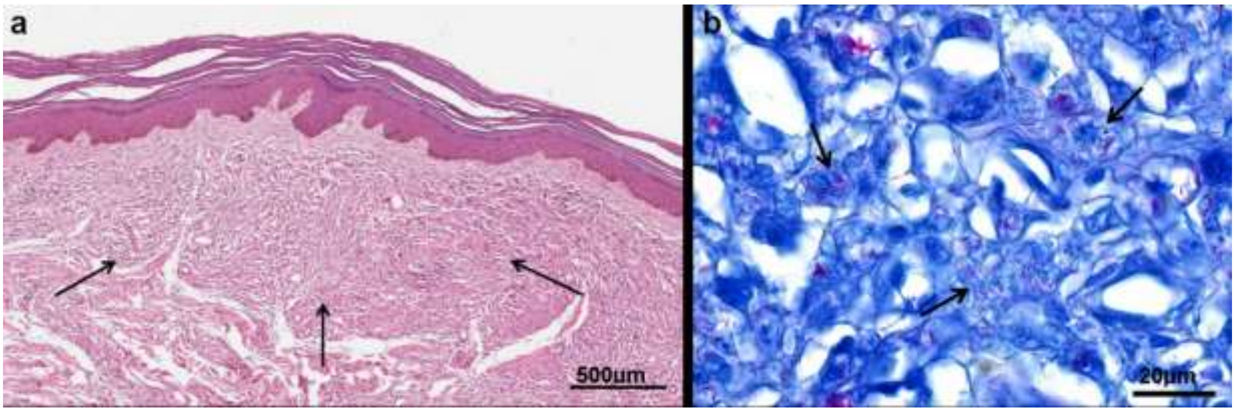


**Extended Data Fig. 5. Histopathology report of skin sample from Zora. a**, Lepromatous leprosy, skin with diffuse histiocytic infiltrate in the dermis. H&E stain, scale bar 500µm **b**, Lepromatous leprosy, skin, acid-fast bacilli in histiocytes. The inflammatory infiltrate consists predominantly of histiocytes admixed with fewer lymphocytes. Histiocytes show foamy or vacuolated cytoplasm and containing bacteria surrounded by a clear zone. Fite-Faraco stain, scale bar 20µm.

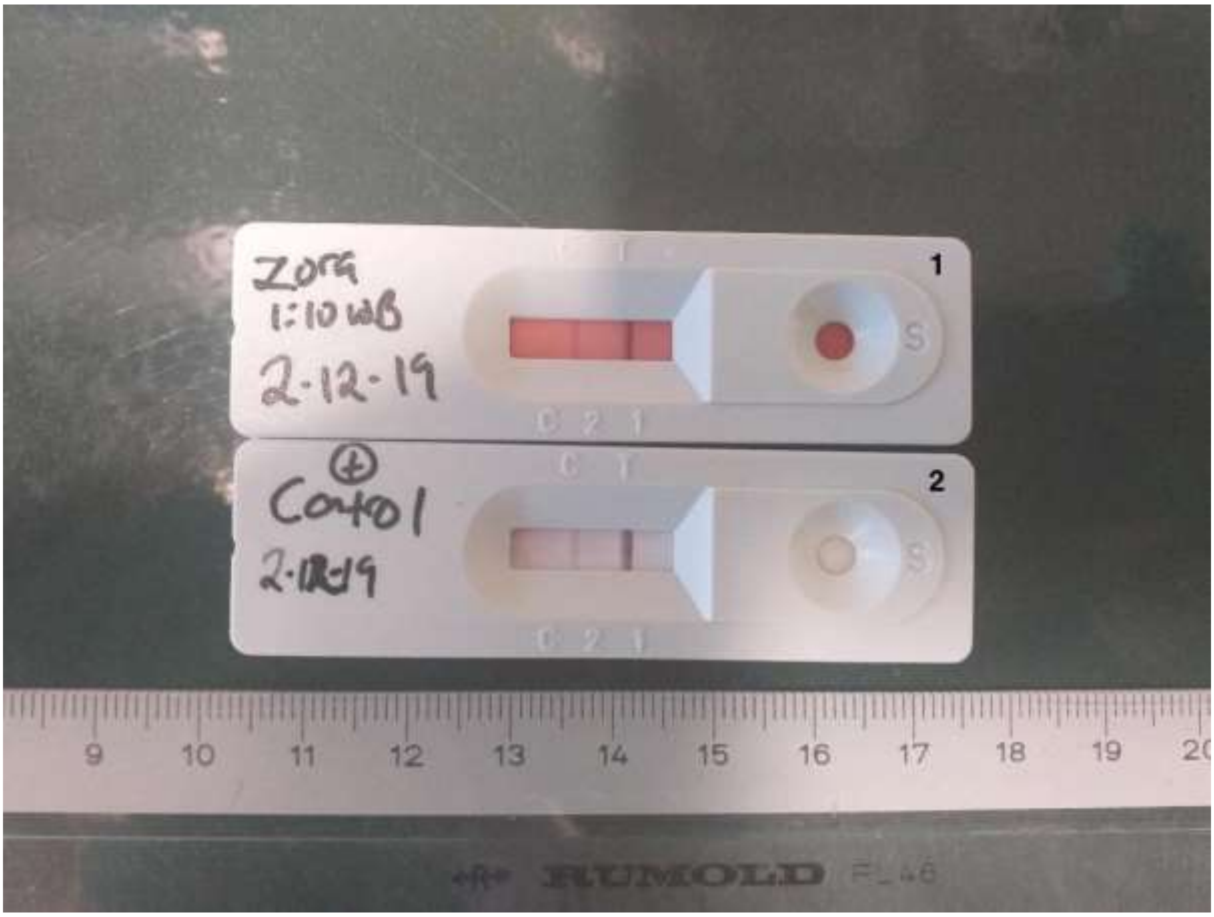


**Extended Data Fig. 6. Lateral flow test result for Zora** whole-blood (1) and the positive control (2). C= control lane; T= test lane.

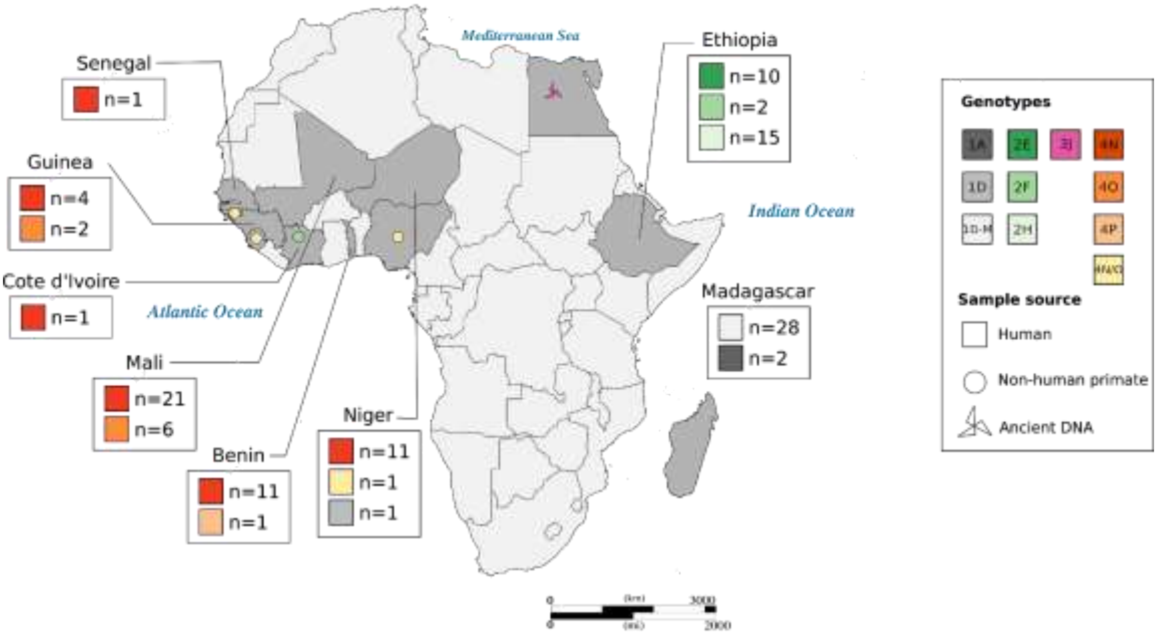


**Extended Data Fig. 7. Geographical distribution of *M. leprae* genotypes in Africa based on genome data.** The genotype 2F has never been reported in West Africa and is the least identified in Ethiopia. The genotype 4N/O was only reported in one human sample from West Africa. Data included only *M. leprae* genomes (Supplementary information Note 5; Supplementary Table 6; Schuenemann et al 2018). The map was downloaded from https://www.amcharts.com/svg-maps/ under a free licence and modified for the current figure in Inkscape, an open source digital illustration software package (https://inkscape.org).

**Extended Data Table 1.** The camera trap (CT) study periods with the focal chimpanzee community within Cantanhez National Park. The number of distinct CT locations for that study period is included (total number of CT locations = 211). Certain CT locations were used in more than one study period. For targeted CT placement if no chimpanzees were filmed for a certain period CTs were repositioned; hence not all cameras were working at the same time. The placement design of CT's was targeted or systematic. Targeted CTs were deployed to maximise detection of chimpanzees (e.g. chimpanzee drumming sites, fruiting trees and trails). Systematic CTs were placed following a survey design maximising independence between CT sites and chimpanzee detection. The CT mode was either set to Photo or Video or both (hybrid) and CTs were active for 24h per day. The start and end dates of each study period are included as well as the number of CT days. CT days are the sum of number of days for each active CT after removing days when cameras were inactive due to malfunctioning, batteries running out, trees falling in front of the CT or theft (total CT days = 28993). The researcher initials are included (JB: Joana Bessa; EB: Elena Bersacola; MR: Marina Ramon).

| Study period | Chimpanzee community | No. CT locations | CT placement | Mode | Start date | End date | CT days | Researcher |
|---|---|---|---|---|---|---|---|---|
| 1 | Caiquene-Cadique Lautchande (2) | 21 | Targeted | Hybrid | 13.09.15 | 16.12.15 | 984 | JB EB |
| 2 | Caiquene-Cadique Lautchande Cambeque Madina Cabante Canamine-Cafache Amindara (7) | 63 | Systematic | Photo | 17.10.16 | 01.03.17 | 3237 | EB |
| 3 | Caiquene-Cadique Lautchande Cambeque Madina Cabante Canamine-Cafache (6) | 50 | Systematic | Photo | 03.06.17 | 15.11.17 | 4435 | EB |
| 4 | Caiquene-Cadique (1) | 21 | Systematic | Photo | 09.07.17 | 05.07.18 | 6838 | EB |
| 5 | Caiquene-Cadique Lautchande Cambeque Madina (4) | 52 | Targeted | Video | 20.02.17 | 03.07.18 | 8023 | JB |
| 6 | Caiquene-Cadique Lautchande Cambeque Madina Cabante Guiledje (6) | 86 | Targeted & Systematic | Video | 04.07.18 | 14.04.19 | 5476 | MR JB |

**Extended Data Table 2.** Individual microsatellite profiles for GB-CC064 and GB-CC068. Alleles are shaded where profile is distinct to that sample.

| | GB-CC064 | | GB-CC068 | |
|---|---|---|---|---|
| **Locus** | **Allele 1** | **Allele 2** | **Allele 1** | **Allele 2** |
| D5s1457 | 111 | 115 | 111 | 115 |
| D13s159 | 161 | 161 | 169 | 171 |
| D2s1326 | 226 | 226 | 226 | 234 |
| D10s1432 | NA | NA | 167 | 167 |
| D16s2624 | 124 | 132 | 116 | 128 |
| D1s207 | 154 | 156 | 154 | 158 |
| D14s306 | NA | NA | 208 | 216 |
| DYs439 | NA | NA | NA | NA |
| D6s311 | NA | NA | 212 | 212 |
| D4s1627 | NA | NA | 225 | 225 |
| HUMFIBRA | 195 | 195 | 191 | 195 |
| amelogenin* | *female* | | *female* | |

**Extended Data Table 3**. Composition of the three multiplex PCRs used to genotype chimpanzee faecal DNA samples GB-CC064 and GB-CC068. All primers at a concentration of 0.2 μM. Size ranges provided for the alleles obtained from two chimpanzees.

| **Multiplex PCR** | **Locus** | **Primers 5' – 3'** | **Dye** | **Product size (bp)** | **Annealing temperature (°C)** |
|---|---|---|---|---|---|
| One | **D5s1457** | Fwd: TGAGGCAATTTGTTACAGAGC<br>Rev: TGCTTGGCACACTTCAGG | FAM | 111-115 | 64 |
| | **D13s159** | Fwd: AGGCTGTGACTTTTAGGCCA<br>Rev: CCAGGCCACTTTTGATCTGT | FAM | 161-171 | |
| | **D2s1326** | Fwd: AGACAGTCAAGAATAACTGCCC<br>Rev: CTGTGGCTCAAAAGCTGAAT | NED | 226-234 | |
| Two | **D16s2624** | Fwd: TGAGGCAATTTGTTACAGAGC<br>Rev: TAATGTACCTGGTACCAAAAACA | NED | 116-132 | Touchdown: 62.5 – 55 |
| | **D1s207** | Fwd: CACTTCTCCTTGAATCGCTT<br>Rev: GCAAGTCCTGTTCCAAGTCT | PET | 140-158 | |
| | **D10s1432** | Fwd: CAGTGGACACTAAACACAATCC<br>Rev: TAGATTATCTAAATGGTGGATTTCC | VIC | 167 | |
| | **D14s306** | Fwd: AAAGCTACATCCAAATTAGGTAGG<br>Rev: TGACAAAGAAACTAAAATGTCCC | PET | 208-216 | |
| | **DYs439** | Fwd: TCCTGAATGGTACTTCCTAGGTTT<br>Rev: GCCTGGCTTGGAATTCTTTT | PET | NA | |
| Three | **Amelogenin** | Fwd: CCTGGGCTCTGTAAAGAATAGTG<br>Rev: ATCAGAGCTTAAACTGGGAAGCTG | VIC | NA* | 59.5 |
| | **HUMFIBRA** | Fwd: GCCCCATAGGTTTTGAACTCA<br>Rev: TGATTTGTCTGTAATTGCCAGC | NED | 191-195 | |
| | **D6s311** | Fwd: ATGTCCTCATTGGTGTTGTG<br>Rev: GATTCAGAGCCCAGGAAGAT | FAM | 212-220 | |
| | **D4s1627** | Fwd: AGCATTAGCATTTGTCCTGG<br>Rev: GACTAACCTGACTCCCCCTC | VIC | 225 | |

* molecular sexing marker.

**Extended Data Table 4. Sequencing data for nonhuman and human primate samples.** S: singleplex; nb: number SD: standard deviation; NR: not relevant, Skin B.: skin biopsy; C.: Chimpanzee; H.: Human

| Library name | Individual | Organ /host | Pool nb | Alignment rate (%) | Mapped reads | Reads used* | Mean fold coverage † | SD † | Genome covered one time (%)‡ | Biosample code |
|---|---|---|---|---|---|---|---|---|---|---|
| TNP418_Li | Zora | Liver / C. | s | 35.1 | 704458 | 10845 | 0.5 | 1.1 | 28.4 | SAMN16207297 |
| TNP418_S1 | Zora | Spleen / C. | s | 92.9 | 24062326 | 276105 | 12.6 | 21.5 | 23.59 | SAMN16207295 |
| TNP418_S2 | Zora | Spleen / C. | s | 91.9 | 25716529 | 294519 | 13.4 | 22.7 | 98.08 | SAMN16207295 |
| TNP418_Lu | Zora | Lung / C. | s | 15.8 | 246681 | 5877 | 0.3 | 0.7 | 17.9 | SAMN16207296 |
| TNP418_merged | Zora | NR | NR | 88.3 | 25267818 | 294922 | 25.8 | 43.8 | **99.7** | - |
| TNP566 | Woodstock | Faecal / C. | s | 18.2 | 1781271 | 25003 | 1.12 | 31.1 | 32.9 | SAMN16207294 |
| CIVW05-07 | Woodstock | Faecal / C. | 2 | 1.1 | 143466 | 4932 | 0.1 | 2.3 | 5.5 | SAMN16207288 |
| CIVW05-08 | Woodstock | Faecal / C. | s | 1.6 | 1075064 | 12775 | 0.3 | 3.1 | 16.9 | SAMN16207289 |
| CIVW11-08 | Woodstock | Faecal / C. | 2 | 0.8 | 118648 | 5166 | 0.1 | 2.5 | 5.1 | SAMN16207290 |
| CIVW28-07 | Woodstock | Faecal / C. | 2 | 1.0 | 138103 | 6632 | 0.1 | 2.6 | 7.7 | SAMN16207291 |
| CIVW28-08 | Woodstock | Faecal / C. | s | 1.4 | 634572 | 8041 | 0.2 | 3.3 | 7.0 | SAMN16207292 |
| CIVW29-08 | Woodstock | Faecal / C. | 2 | 1.0 | 160326 | 4571 | 0.1 | 2.5 | 3.5 | SAMN16207293 |
| CIVW_all | Woodstock | NR | NR | 1.3 | 2270179 | 26883 | 0.6 | 3.8 | 36.2 | - |
| GB-CC064_1 | NR | Faecal / C. | s | 2.4 | 1161036 | 890751 | 6.3 | 4.9 | 97.9 | - |
| GB-CC064_2 | NR | Faecal / C. | s | 1.4 | 2619804 | 667842 | 31.6 | 111.7 | 99.9 | - |
| GB-CC064_merged | NR | NR | NR | 2.2 | 3025429 | 595745 | 39.3 | 60.2 | 99.9 | SAMN16207298 |
| Bn10-71 | NR | Skin B. /H. | s | 86.6 | 1200600 | 1025025 | 23.4 | 13.1 | 99.9 | SAMN16207303 |
| Bn7-37 | NR | Skin B. /H. | s | 55.9 | 6887990 | 3237463 | 99.9 | 15.5 | 100 | SAMN16207301 |
| Bn7-38 | NR | Skin B. /H. | s | 51.0 | 20317528 | 5510687 | 170.1 | 10.9 | 99.9 | SAMN16207302 |
| Bn9-59 | NR | Skin B. /H. | s | 67.0 | 1898621 | 1440783 | 32.9 | 18.1 | 100 | SAMN16207304 |
| Bn9-66 | NR | Skin B. /H. | NR | 99.1 | 11680598 | 4530817 | 103.7 | 25.4 | 100 | SAMN16207305 |
| Bn9-67 | NR | Skin B. /H. | NR | 97.9 | 11875974 | 4659249 | 106.7 | 23.7 | 100 | SAMN16207306 |
| CI-7 | NR | Skin B. /H. | NR | 48.9 | 21847244 | 204700 | 4.7 | 5.7 | 98.3 | SAMN16207307 |
| ML11-101 | NR | Skin B. /H. | NR | 55.5 | 4231957 | 1343804 | 61.2 | 8.6 | 100 | SAMN16207308 |
| Ml11-102 | NR | Skin B. /H. | NR | 78.8 | 4087000 | 2014082 | 93.7 | 8.6 | 100 | SAMN16207309 |
| Ml11-103 | NR | Skin B. /H. | NR | 41.0 | 2977125 | 1876824 | 86.1 | 9.2 | 100 | SAMN16207310 |
| Ml12-109 | NR | Skin B. /H. | NR | 98.0 | 13586511 | 1678386 | 38.5 | 11.6 | 100 | SAMN16207311 |
| Ml12-110 | NR | Skin B. /H. | NR | 96.2 | 3622404 | 986042 | 22.6 | 8.0 | 100 | SAMN16207312 |
| Ml2-7 | NR | Skin B. /H. | NR | 99.2 | 7734485 | 4264605 | 129.1 | 9.5 | 100 | SAMN16207313 |
| Ml3-17 | NR | Skin B. /H. | NR | 98.3 | 7354925 | 4164480 | 127.8 | 9.6 | 100 | SAMN16207314 |
| Ml6-52 | NR | Skin B. /H. | NR | 91.3 | 3716990 | 2715341 | 82.8 | 8.3 | 100 | SAMN16207315 |
| Ng19-42 | NR | Skin B. /H. | NR | 4.7 | 768700 | 620403 | 29.2 | 5.7 | 100 | SAMN16207316 |
| Ng22-45 | NR | Skin B. /H. | NR | 1.6 | 282398 | 250477 | 11.6 | 3.6 | 99.9 | SAMN16207317 |
| Ng27-58 | NR | Skin B. /H. | NR | 89.5 | 4610071 | 2865550 | 65.5 | 25.1 | 100 | SAMN16207318 |
| Ng27-60 | NR | Skin B. /H. | NR | 64.8 | 1673215 | 1348408 | 30.8 | 14.4 | 100 | SAMN16207319 |

| | | | | | | | | | | |
|---|---|---|---|---|---|---|---|---|---|---|
| Ng29-64 | NR | Skin B. /H. | NR | 76.1 | 1430979 | 1170655 | 26.8 | 13.9 | 99.9 | SAMN16207320 |
| Sen1-1 | NR | Skin B. /H. | NR | 91.6 | 1560006 | 1289143 | 29.5 | 14.6 | 100 | SAMN16207321 |

∗ after quality check removing duplicates † no duplicates ‡ at least

**Extended Data Table 5. Primers used for the identification of *M. leprae* in chimpanzee tissues and faeces, diet analysis, the genotyping of *M. leprae* strains and confirmation of chimpanzee origin of the samples. Fwd: forward; Rev: reverse**

| PCR system and target | Primer pair 5´-3´ | Product size (bp) | Annealing temperature (°C) |
|---|---|---|---|
| RLEP_Primary PCR | Fwd: TGCATGTCATGGCCTTGAGG<br>Rev: CACCGATACCAGCGGCAGAA | 129 | 58 |
| RLEP_Nested PCR | Fwd:TGAGGTGTCGGCGTGGTC<br>Rev: CAGAAATGGTGCAAGGGA | 99 | 58 |
| Fusion_M13_RLEP PCR | Fwd: GTAAAACGACGGCCAGTGAGGTGTCGGCGTGGTC<br>Rev: CAGGAAACAGCTATGACCAGAAATGGTGCAAGGGA | 139 | 58 |
| 18kDA _Primary PCR | Fwd: TCATAGATGCCTAATCGACTG<br>Rev: GGCACATCTGCGGCCAGCA | 136 | 55 |
| 18kDA _Nested PCR | Fwd: ATCGACTGTTGTTTGCGCAAC<br>Rev: CCAGCAACCGAAATGTTCGGA | 110 | 55 |
| Fusion_M13_18kDA PCR | Fwd:GTAAAACGACGGCCAGATCGACTGTTGTTTGCGCAAC<br>Rev: CAGGAAACAGCTATGACCCAGCAACCGAAATGTTCGGA | 150 | 55 |
| 16S Mammal identification PCR (diet analysis) | 16Smam1: CGGTTGGGGTGACCTCGGA<br>16Smam2: GCTGTTATCCCTAGGGTAACT<br>16Smam_Human_blocker: CGGTTGGGGCGACCTCGGAGCAGAACCC<br>16Smam_Pig_blocker: CGGTTGGGGTGACCTCGGAGTACAAAAAAC | 130 | 64 |
| Mammal identification fusion PCR (diet analysis) | 16Smam1_Illumina_adapter: TCGTCGGCAGCGTCAGATGTGTATAAGAGACAGCGGTTGGGGTGACCTCGG<br>16Smam2_Illumina_adapter: GTCTCGTGGGCTCGGAGATGTGTATAAGAGACAGGCTGTTATCCCTAGGGTAACT | 160 | 64 |
| 16S Species confirmation PCR | 16Smam1: CGGTTGGGGTGACCTCGGA<br>16Smam4: AGATAGAAACCGACCTGGAT | 300 | 64 |
| Genotyping of leprosy strain infecting Woodstock | ml0048-Fwd*: ATACCGTGACGCGGATAAAC<br>ml0048-Rev*: GTAGCCAGTCCAAGGCAATC<br>ml0565-Fwd**: AGCTGAGGTTGACCTGGAA<br>ml0565-Rev**: GTAGATTGGCGTCGTCAAAA | 576<br>561 | 55<br>57 |

*Mutation C1193T in *ml0048* (genome position 60123); a T is found in TNP418 and TNP566
**Mutation C319T in *ml0565* (genome position 683097); a T is found in TNP418 and TNP566
