## Supplementary Information for "Leprosy in wild chimpanzees"

**Description of Content:**

This file contains the supplementary information on chimpanzee socioecology, and supplementary methods and results for each of the observational, molecular, genomic, histopathological and serological investigations. We also include five Supplementary Videos with titles.

**Note 1: Chimpanzee Socioecology**

Chimpanzees are semi-terrestrial, diurnal, and social. They live in mixed-sex groups called communities. Neighbouring communities use discrete territories and adult males monitor overlapping boundary zones[68]. Chimpanzees have a high degree of fission-fusion dynamics, whereby individuals of a community split into small subgroups or parties which are scattered throughout the home range, with party size and composition changing frequently[69]. Their society is male-dominated and patrilineal. Females normally emigrate to a neighbouring community when they reach puberty, whereas males remain in their natal group.

The natural diet of chimpanzees is broadly omnivorous, comprising diverse plant and animal foods, although chimpanzees preferentially seek out ripe fleshy fruits[70]. In agroforest landscapes, chimpanzees often maintain a diet dominated by ripe fruit by incorporating human crops[45,71]. Chimpanzee communities exhibit variation in their diets, including their consumption of prey species. Common prey taxa include rodents and ungulates, as well as other nonhuman primates such as colobine monkeys (*Piliocolobus* and *Colobus* spp.), baboons (*Papio* spp.), guenons (*Chlorocebus* spp.) and galagos (*Galago senegalensis*)[72]. Chimpanzees use nests for sleeping at night and resting during the day. Normally they build a new arboreal nest for sleeping every night, but sometimes existing nests are reused[73].

### Note 2: Monitoring of individuals and longitudinal observation at the study sites

#### Observations of leprosy like-lesions in chimpanzee from CNP

We conducted 28,993 camera-trapping days from 13th September 2015 to 14th April 2019 (Extended Data Table 1). We deployed camera traps to survey eight putative chimpanzee communities at 211 locations (Supplementary Fig. 1b), covering approximately 320km$^2$ of CNP. We organised images containing chimpanzees into independent events and 60 minutes was used as the minimum time interval separating independent events[74–76]. Some chimpanzees displayed physical characteristics that might be attributed to early leprosy (e.g. skin hypopigmentation, hair loss); however, as leprosy could not be confirmed, these cases were noted but not included in analyses. We gave chimpanzees showing advanced leprosy-like symptoms an ID to enable future identification. Images and videos were carefully reviewed and data files including chimpanzees showing leprosy-like lesions were analysed to identify and monitor progression of symptoms over time.

**Terms used for Clinical Manifestations of Leprosy in Chimpanzees**

**Callosity:** Thickening and hardening area of the skin.

**Claw hand:** Hyperextension of metacarpophalangeal joint, flexion of proximal and distal interphalangeal joints of the $4^{th}$ and $5^{th}$ fingers. Typical clinical sign of human leprosy cases.

**Hypopigmentation:** lightening of the skin/ loss of skin colour.

**Nodule:** A circumscribed elevated, solid skin lesion.

**Patch:** The skin appears dry with hair loss but is not visibly raised or scaly. Patch is not defined based on the size of the affected area as in combination with the absence of direct clinical inspection of individuals; hair cover can impede the identification of small patches. The term patch is used if the quality of the video or photo does not allow us to confirm the affected area is a plaque.

**Plaque:** A broad raised area on the skin that appears scaly and dry, with hair loss in affected area.

**Ulceration:** Lesion that cracks and bleeds, forming an open wound.

### Descriptions of Individual Disease Progression

### Cantanhez National Park (CNP), Guinea-Bissau

*Rita (Caiquene-Cadique; images from 2013 to 2020)*

2013: *Head:* asymmetrical thickening of ear margins and brow ridge; hypopigmentation of upper lip and nose; small nodule on the lower lip and left ear. *Body:* no visible hair loss or skin patches. Generally good physical condition.

2015: *Head:* new cluster of larger nodules between the upper lip and nose; multiple small nodules on the eyelids, cheek, ears margins, lower lip, and brow ridge. *Body*: small patches on wrists, knees and elbows, with some hair loss. Unable to confirm clinical signs on hands and feet due to muddy terrain in mangroves.

2016: *Head:* nodules increase in size and appear more prominent on lower lip and brow ridge; thinning of hair/ hair loss. *Body:* plaques on the joints including wrists, knees and elbows; thinning of hair/ hair loss on limbs; apparent swelling of feet and hands (unable to confirm due to muddy terrain in mangroves).

2017: *Head:* nodules increase in number, apparent swelling and reddening of nodules; increase in hair thinning; facial disfigurement. *Body:* increase in hair thinning; scaly, plaques in several areas including the shoulder, lower back, knee, elbow; disfigurement of feet and hands including claw hand (Supplementary Video 1).

2018: *Head:* face and ears are completely covered by larger nodules that appear shiny; facial disfigurement. *Body:* generalised hair loss on limbs and lower back; nodules on hands and feet; claw hand more severe (Supplementary Videos 2 and 3); nodulation and swelling of fingers and toes, which are becoming disfigured; some plaques are ulcerated; weight loss.

*2020: Head:* face becomes slightly ulcerated; ear margin looks thinner with scarring, and eyelids and brow ridge look smoother. *Body:* continuous hair loss on the limbs and lower back;

plaques on wrists, upper arm, legs and lower back are larger and more ulcerated; increased swelling of feet and hands.

*Brinkos (Caiquene-Cadique; images from 2015 to 2020)*

2015: *Head*: skin thickening, multiple large nodules affecting face (cheek, chin, eyelids, lips) and brow ridge, nodules visible on left ear. *Body*: Large areas of the lower back, and upper and lower limbs covered by plaques; hair loss on lower back, shoulder, elbow, upper arm, and upper leg (left side visible). Unable to confirm clinical signs on hands and feet due to muddy terrain in mangroves.

2016: *Head*: nodules on and around nose and upper lip increase in size; apparent swelling and reddening of nodules, with some on the left ear and brow ridge becoming ulcerated; large nodules on visible left ear margin; facial disfigurement. *Body*: presence of plaques as in 2015; hair loss affecting vast areas of the lower back and limbs.

2017: *Head*: thinning of hair on frontal region of the head; larger nodules completely covering face; nodules on ears increasing in number and size; ulceration of lesions across the face; facial disfigurement. *Body*: vast plaques covering the lower back, shoulders, arms and legs; hanging nodule in anogenital area; swelling of hands, feet and fingers.

*2018*: *Head*: hair loss, facial disfigurement, ulceration of nodules. Larger nodules form on ears with a hanging large nodule on both earlobes, and a hanging lower lip. *Body*: ulcerated hands and feet, swelling of fingers, nodules on nipples. The plaques appear larger and thicker and affect the sides of the body, with some ulcerated; increase hair loss.

2020: *Head*: Increased ulceration of nodules. *Body*: plaques cover the body and present multiple crusts and scarring; swelling of hands, feet and fingers; more extensive hair loss (hair now only present on the back of the head and neck, the inner parts of upper and lower limbs, and the central area of the back and abdomen).

*Jimi (Lautchande; images from 2015-2018)*

2015: *Head:* General swelling of face, especially cheek and brow ridge, and hypopigmented spots on brow ridge (one) and below nostrils. Ears do not appear affected. *Body:* general good condition. A small patch of hair loss can be observed on right wrist and on right knee. Hands and feet not visible.

2016: *Head:* ear margins and brow ridge swelling, small nodules are visible on upper lip, and brow ridge; scaly brow ridge and ear margins. *Body:* small plaque on upper back; buttocks, hands, and feet present nodules.

2017: *Head:* completely covered by multiple nodules, which are shiny and reddish, some are ulcerated; thickening of ear margins. *Body:* new and larger plaques appear on other parts of the body including the wrists, lower limbs, lower back, buttocks and testicles; hands and feet present nodules that appear swollen and ulcerated; calloused skin on the bottom of the feet.

2018: *Head:* as in 2017. *Body:* plaques on the lower back become larger and drier and new plaques appear on the upper area, following vertical strip patterns.

*Baaba (Cambeque; images available for 2017 and 2018)*

2017: *Head:* multiple hypopigmented nodules on brow ridge, cheek, upper and lower lip; ears with thick margins and nodules; hair thinning. *Body:* multiple small plaques present on the upper and lower limbs, back, abdomen and shoulders.

2018: *Head:* nodules located between the upper lip and nostrils become larger and redder. *Body:* increase in hair loss/thinning; hands and feet not visible.

**Tai National Park, Côte d'Ivoire**

*Woodstock (from 2018/06 to 2020/07)*

2018/06: *Head*: First hypopigmented nodular lesions appear all over the face: eye orbits (1 large nodule under right eye), eyelids (multiple small nodules on left eyelid, and one on the right eyelid), brow ridges thickened (multiple smaller and larger nodules, one very large prominent in the middle of the left brow), earlobe (1 large nodule on the left earlobe), lips (one large nodule on lower left side), nose (one small nodule under nasal septum), one small nodule on the left side of face between the lips and the nose.

*Body*: Swelling and hypopigmentation of both hands. Ulceration of the skin at the level of the proximal interphalangeal joint on the 4$^{th}$ finger of the right hand. No apparent swelling, hypopigmentation, or lesions on the feet.

*Behaviour*: No apparent social, feeding, or locomotive behavioural change. Woodstock does not scratch or groom the lesions. Other individuals of the group do not appear interested in the lesions.

2018/10:

*Head*: The existing nodules increase in size; multiple new small ones appear on the areas around and under the nose on both sides of the face, as well as on the eyelid margins and brow ridges. Development of a mucopurulent discharge from the left eye, lower eyelid turns outward. Left ear appears swollen, ear margins thickened, as the nodule continues to grow.

*Body*: Hair loss and ulceration on the dorsal part of right wrist. Ulceration of the skin at the level of metacarpophalangeal and proximal interphalangeal joints on the 2$^{nd}$ finger of the right hand. On the left hand there are no apparent lesions or ulcerations, patches on the wrist can be observed. Slight hypopigmentation visible on both feet, but no other lesions or swelling apparent.

*Behaviour*: No apparent social or feeding behavioural change. Due to ulcerations on the right knuckles, Woodstock occasionally walks on his right wrist instead of knuckles, causing a slight limp, although this does not impact his walking speed. Other individuals from the group groom the ulcerations but not the lesions with intact skin.

2019/01:

*Head*: Majority of existing nodules increase in size and become pedunculated, nodule under eye shrinks. New small ones appear on the lips. Both ears swollen, ear margins thickened and disfigured, the left ear nodule continues to grow.

*Body*: Hands are swollen and hypopigmented, no ulceration visible. Both feet are swollen and hypopigmented; ulcerated plaques appear on the lateral side of the left foot.

*Behaviour*: Woodstock appears to have difficulty moving, with no apparent changes in social or feeding behaviour.

2019/04:

*Head*: The existing nodules increase in size, nodules located close to each other fuse together. Both ears are swollen, ear margins thickened, the left earlobe is completely disfigured; the right ear starts to become disfigured. White discharge from left eye still present. New nodule formation on chin, nodules on lips become pedunculated. Suspected start of nasal involvement.

*Body*: Both hands are slightly swollen and hypopigmented. Small skin ulceration appears at the proximal interphalangeal joint on the 2$^{nd}$ finger of the right hand. On the 4$^{th}$ finger of the left hand, loss of nail plate can be observed, nail bed becomes exposed; on the 3$^{rd}$ and 5$^{th}$ fingers, early stage of abnormal nail overgrowth can be seen. The feet do not appear swollen, only hypopigmented, the plaques on the left foot are present without any ulceration. Degradation of nails on the left foot can be observed. Hypopigmentation of scrotum.

*Behaviour*: Woodstock left the community for three weeks. On his return, he displayed no apparent change in behaviour, although he appears to have lost weight.

2019/09:

*Head*: Facial lesions increase in size; many new ones on brow ridge and above the lips become pedunculated.

*Body*: On the 1st and 2nd fingers of the left hand, loss of nail plate can be observed, nail bed becomes exposed.

*Behaviour*: Woodstock left the community for three months. On his return he appeared to be behaving normally, although his movement was slower, and he was biting his fingernails (Supplementary Video 4).

2020/04:

*Head*: Lesions on brow ridge merge together. Nodules appear smaller than before.

*Body*: No visible abnormalities on the right hand. On the left hand, the nail of the 4th finger shows an advanced stage, the 3rd and 5th fingernails show early stage of abnormal nail overgrowth. Hypopigmentation and nail involvement on the right and left feet can be observed. On the 1st and 4th toenails of the left foot complete, and on the 3rd toenail, partial loss of nail plate can be observed.

*Behaviour*: Woodstock displays normal social behaviour, although his body condition is worse, and he is moving more slowly.

2020/07:

*Head*: Brow ridge protruding. Both ears swollen, ear margins thickened and disfigured. Nasal involvement becomes apparent.

*Body*: Body condition slightly worse. Both hands are swollen and hypopigmented. Skin ulcerations present on the level of $3^{rd}$, $4^{th}$ and $5^{th}$ proximal interphalangeal joints of the right hand, as well as on the level of the $3^{rd}$, $4^{th}$ and $5^{th}$ proximal interphalangeal joints of the left hand. Advanced stage of abnormal nail overgrowth, of the $4^{th}$ and $5^{th}$ fingernail of the left hand. Possible observation of "claw hand" on left hand (Supplementary Video 5). Reddening and ulceration of scrotum; fresh blood could be seen.

*Behaviour*: Woodstock displays no apparent change in feeding and social behaviour. He occasionally walks on the wrist instead of knuckles, and sometimes moves slower than other individuals. He touches and licks ocular discharge, and occasionally licks his finger ulcers, but does not pay attention to ulcers on his scrotum.

*Zora (from 2001-2009)*

Retrospective PCR screening (see Note 4.1) revealed presence of *M. leprae* DNA in faecal and necropsy samples of another individual, Zora. The individual was estimated to be 57 years-old at the time of her death in 2009. From molecular investigations at the time, Zora was diagnosed with a tuberculosis infection and viable *M. tuberculosis* bacilli were isolated from mesenteric lymph nodes and sequenced[77].

Archived photos of Zora from 2001-2009 were reviewed for leprosy like lesions. They reveal marked hypo-pigmented nodular lesions on the ear pinnae, eyebrows and a few lesions on lips, maxillofacial area and distal extremities (Extended Data Fig 4). However, even though this individual was infected since 2002 based on the earliest detection of *M. leprae* in the faeces, it was difficult to reliably determine when the symptoms of leprosy appeared. In old female chimpanzees, hypopigmentation is a natural phenomenon and too few archived photos were available to thoroughly assess disease progression.

**Note 3: Diet analysis and genetic characterisation**

**Diet analysis in chimpanzees at CNP**

Molecular identification of mammalian prey DNA in 121 chimpanzee faeces from CNP was performed using a metabarcoding approach, as previously described[58]. Amplicon deep sequencing was conducted on PCR products generated by targeting a 130 bp fragment of the 16S rRNA in presence of blocking primers (human and swine). A custom pipeline using OBITools v1.1.8 was used for downstream taxonomic assignment of each read to the family, genus and order level[78]. To be included, greater than 10 reads needed to be assigned to the prey species and at least 0.1% of assigned amplicon reads assigned to the prey species. Of the 118 faeces analysed, diet outcomes were three Campbell's monkeys (*Cercopithecus campbelli*) (GB-L001, GB-CB007, GB-CB016), one West African large spotted genet (*Genetta pardina*) GB-CC035), and one red river hog (*Potamochoerus porcus*) (GB-CC070).

**Genetic characterisation confirming chimpanzee origin at CNP**

At CNP chimpanzees are not habituated. Therefore, defecation is not normally observed. Instead, chimpanzee faeces are collected under nests or in areas where other signs of recent chimpanzee presence are detected. While chimpanzee faeces are easy to recognise, it cannot be ruled out that on rare occasions faeces from another species are accidentally collected. To confirm that samples containing *M. leprae* DNA or mammal DNA in diet analyses were indeed of chimpanzee origin, we performed a mammal PCR targeting a larger fragment (300 bp) of the mitochondrial 16S rRNA gene without using blocking primers. As this is a PCR we regularly use, we included a Uracil-DNA glycosylase (UNG) step to avoid contamination with PCR products. PCRs were performed in 25µL reactions including 200ng of DNA, 1.25U of high-fidelity Platinum Taq™ polymerase, 10x PCR buffer, 200µM dNUTPs, 4mM MgCl2, and 200nM of both forward and reverse primers (Extended Data Table 5). The thermal cycling conditions were as follows; 45°C for 7 min (UNG treatment), 95°C for 5 min (UNG inactivation), 95°C for 5 min (denaturation), followed by 42 cycles of 95°C for 30 sec, 64°C for 30 sec, and 72°C for 1 min, and a final elongation step at 72°C for 10 min. PCR products were visualised on a 1.5% agarose gel stained with GelRed®, bands were purified using the Purelink Gel extraction kit, and sequenced after Sanger's method using forward and reverse primers. Cleaned sequences were compared to publicly available nucleotide sequences using the Basic Local Alignment Search Tool (BLAST)[56]. All faecal samples from CNP positive for *M. leprae* or mammalian DNA originated from chimpanzees.

**Note 4: Genetic identification of samples from infected chimpanzees at CNP**

DNA samples were quantified and checked for quality using a DropSense96 (Trinean) spectrophotometer and a Qubit (Thermo Fisher Scientific, MA, USA) fluorometer (dsDNA BR Assay Kit). DNA concentration of all three samples was found to be less than 1ng/µl, to increase the quantity of DNA available for downstream applications all samples were purified and concentrated using a QIAquick PCR Purification Kit (QIAGEN, Germany). Extractions were subsequently used at working concentrations of 2ng/µl.

Multiplex compositions, fluorescent dye choices, and PCR cycling conditions were obtained[55]. Microsatellite loci were amplified in 15 µl volumes of a PCR mix consisting of 1x QIAGEN Multiplex PCR Master Mix®, primers at a concentration of 0.2µM, (Extended Data Table 3), and approximately 8-10ng of DNA. In total 11 loci were targeted and one sexing marker, these are identified as multiplex reactions M1, M2, and M3[55]. Reactions were carried out in Applied Biosystems Veriti™ Thermal Cyclers. PCR products were run on a 3500 Applied Biosystems® sequencer and included the size standard GeneScan™ 600 LIZ™ dye (Thermo Fisher Scientific, MA, USA). Fragment analysis was performed using Geneious® 9.1.5[79]. Allele scoring was semi-automated, with anticipated bins and peak calling automated, followed by visual inspection to avoid scoring errors and interpretation of low signal intensity peaks.

**Note 5: Whole genome sequencing of *M. leprae* strains infecting chimpanzees and humans**

**Post-mapping analysis and geographical origin of strains**

A total of 99.7 and 99.9% of the *M. leprae* genome were recovered from tissue samples of Zora (Côte d'Ivoire) and from the faecal sample GB-CC064 (Guinea-Bissau) with a mean coverage of 25.8X and 39.3X, respectively (Extended Data Table 4). For Woodstock, 23.5% of the *M. leprae* genome was recovered from faecal samples with a mean coverage from 1.1X (Extended Data Table 4). This incomplete genome was not included in the phylogenetic analysis.

Besides, 21 genomes from five West African countries (Niger, Mali, Benin, Côte d'Ivoire and Senegal) were added to the analysis with a coverage ranging from 4.7 to 170-folds (Extended Data Table 4). With the addition of previously sequenced genomes, the current dataset is composed of 64 genomes from West Africa covering six countries (Supplementary Table 6 and Extended Data Fig 7).

**Comparative genomics of *M. leprae* strains from naturally infected non-human primates and humans**

*GB-CC064 from Guinea-Bissau*

Inside the branch 4, GB-CC064 shared a common ancestor with all type 4 genotypes, deriving from common ancestors with the 3M and 3L genotypes, all reported in humans. GB-CC064 does not specifically cluster with any previously sequenced human or animal strains (Fig 3a), but branches off separately in the so-called genotype 4N/O which shares the informative position of 4N and 4O as described by Monot and colleagues (Fig 1b). It is unknown which *M. leprae* strains circulate in humans in Guinea-Bissau, because no human-infecting *M. leprae* strain from this country has ever been sequenced, and no leprosy cases were reported in humans in 2018[5]. While it is unlikely that the rare 4N/O strain plays an important role in humans in

Guinea Bissau, the lack of data makes it impossible to rule out a recent human source of infection for chimpanzees in Guinea-Bissau.

GB-CC064 harboured 23 specific variants when compared to the other strains from the branch 4 including one deletion in a pseudogene and one SNP (G361545C, Pro137Ala) in *ml0279* coding for a thioesterase, shared with Airaku-3, a strain from Japan, belonging to the genotype 1D (Supplementary Table 6).

*M. leprae strains infecting Zora and Woodstock*

Comparative genomics showed that the strain TNP-418 recovered from Zora belongs to the branch 2F currently composed of ancient human strains from medieval Europe and modern human strains from Ethiopia but cluster independently from others (Fig 2b). The genotype 2F has never been described in West Africa and was reported in only two out of 27 samples from Ethiopia, as well as in the Middle East (Turkey and Iran)[59] and in medieval Europe (Denmark, United Kingdom, Sweden and Italy). TNP-418 harboured 34 specific SNPs compared to the other 285 genomes (Supplementary Table 6). The occurrence of these mutations was checked manually on IGV v2.3.79 in sequencing data from Woodstock. Ten positions were covered by one to five reads which showed presence of the same variants found in TNP-418 (Supplementary Table 6), and presence of two additional SNPs was confirmed by PCR. Therefore, it is likely that Woodstock was infected with a similar strain to Zora's (probable transmission within the group or from the same source). Such low genetic diversity is expected based on the data observed in armadillos in the USA, red squirrels from Brownsea Island and in strains from humans living in the same household or village[10,12].

*Dating*

The most recent common ancestor (MRCA) of all the *M. leprae* strains from branch 2F is estimated to be 2772 years old [95% highest posterior density (HPD) of 2261–3267 ya] (Fig 3a). The estimated mean age of the MRCA is about 100 y older compared to the estimates derived from previous reports[27,60]. The shift probably resulted from the addition of TNP-418 since the MRCA for the other branches are similar to the values previously published. The MRCA of TNP-418 is estimated to have diverged from all other 2F strains 1873 ya [95% HPD of 1564–2204 ya], i.e., in the 2$^{nd}$ century C.E (Fig 3a). The probability of distribution calculated by BEAST (posterior = 1) and the bootstraps value from MEGA (bootstrap 76) confirmed the placement of TNP-418 in the overall tree as well as its placement as a unique representative of its own sub-group inside the branch 2F.

The MRCA of all the *M. leprae* strains from branch 4 is estimated to be 1877 years old [95% HPD of 1502–2295 ya] while the MRCA of the genotype 4N/O with the other genotype 4 (4N, 4O and 4P) estimated to be 1437 years old [95% HPD of 1132–1736 ya] (Fig 3a). The MRCA of all 4N/O genotype was circulating during the 7$^{th}$ century (1368 years old, 95% HPD 1074-1668).

### Note 6: Ethical standards

#### Nonhuman primates

All data were collected in accordance with Best Practise Disease and Monitoring Guidelines of the Great Ape Section of IUCN Primate Specialist Group[50]. The collection of samples was strictly non-invasive. All proposed data collection and analyses adhered strictly to ethics guidelines of the Association for the Study of Animal Behaviour (UK). Ethical approval for targeted leprosy camera trap surveys and faecal sample collection at CNP, Guinea-Bissau, was granted by the University of Exeter, UK. The Institute for Biodiversity and Protected Areas (IBAP) in Guinea-Bissau approved and collaborated directly on all aspects of this research. Ethical approval for the work at Tai Chimpanzee Project was given by the Ethics Commission of the Max Planck Society. The Centre Suisse de Recherches Scientifiques en Côte d'Ivoire collaborates on the research at TNP.

#### Human individuals

This study was carried out under the ethical consent of the WHO Global Leprosy Program surveillance network. All subjects gave written informed consent in accordance with the Declaration of Helsinki.

**Supplementary Videos**

**Supplementary Video 1.** Videoclip shows adult female chimpanzee, Rita, with claw hand at CNP (dated April 2017).

**Supplementary Video 2.** Videoclip shows adult female chimpanzee, Rita, at CNP (dated May 2018).

**Supplementary Video 3.** Videoclip shows adult female chimpanzee, Rita, at CNP (dated November 2018).

**Supplementary Video 4.** Videoclip shows adult male chimpanzee, Woodstock, at TNP biting his fingernails (dated April 2019).

**Supplementary Video 5.** Videoclip shows adult male chimpanzee, Woodstock, with claw hand at TNP (dated July 2020).

**SUPPLEMENTARY REFERENCES**


68. Watts, D. & Mitani, J. Boundary patrols and intergroup encounters in wild chimpanzees. *Behaviour* **138**, 299–327 (2001).

69. Nishida, T. The social group of wild chimpanzees in the Mahali Mountains. *Primates* **9**, 167–224 (1968).

70. Goodall, J. *The chimpanzees of Gombe: patterns of behavior*. (Belknap Press of Harvard University Press, 1986).

71. McLennan, M. R. & Hockings, K. J. Wild chimpanzees show group differences in selection of agricultural crops. *Sci. Rep.* **4**, 5956 (2014).

72. Newton-Fisher, N. E. The hunting behavior and carnivory of wild chimpanzees. in *Handbook of Paleoanthropology* (eds. Henke, W. & Tattersall, I.) 1661–1691 (Springer, 2015). doi:10.1007/978-3-642-39979-4_42.

73. Fruth, B. & Hohmann, G. Nest building behavior in the great apes: the great leap forward? in *Great Ape Societies* (eds. Marchant, L. F., Nishida, T. & McGrew, W. C.) 225–240 (Cambridge University Press, 1996). doi:10.1017/CBO9780511752414.019.

74. Garriga, R. M. *et al.* Factors influencing wild chimpanzee (*Pan troglodytes verus*) relative abundance in an agriculture-swamp matrix outside protected areas. *PLoS ONE* **14**, e0215545 (2019).

75. Rovero, F. & Marshall, A. R. Camera trapping photographic rate as an index of density in forest ungulates. *J. Appl. Ecol.* **46**, 1011–1017 (2009).

76. Rovero, F., Martin, E., Rosa, M., Ahumada, J. A. & Spitale, D. Estimating species richness and modelling habitat preferences of tropical forest mammals from camera trap data. *PLoS ONE* **9**, e103300 (2014).

77. Coscolla, M. *et al.* Novel *Mycobacterium tuberculosis* complex isolate from a wild chimpanzee. *Emerg. Infect. Dis.* **19**, 969 (2013).

78. Boyer, F. *et al.* obitools: a unix-inspired software package for DNA metabarcoding. *Mol. Ecol. Resour.* **16**, 176–182 (2016).

79. Kearse, M. *et al.* Geneious Basic: an integrated and extendable desktop software platform for the organization and analysis of sequence data. *Bioinformatics* **28**, 1647–1649 (2012).
