## Supplementary Table Titles for "Leprosy in wild chimpanzees"

**Supplementary Table captions**

Sup Table 1: Chimpanzee DNA samples tested for *M. leprae*.
Sup Table 2: List of the 286 genomes used in this study
Sup Table 3: SNP table for the 286 genomes included in this study
Sup Table 4: BEAST input SNP table
Sup Table 5: Nucleotide positions manually annotated in the SNP table for GB-CC064 and TNP418
Sup Table 6: Autapomorphic SNPs from GB-CC064 and TNP418.
